# Regional cerebral arterial elasticity is associated with cardiovascular health in cognitively healthy older adults

**DOI:** 10.64898/2026.07.30.740814

**Authors:** Jenna E. Johnson, Nicholas Ware, Sarah J. Johnson, Gabriele Gratton, Kathy A. Low, Daniel Barker, Montana Hunter, Monica Fabiani, Ashleigh E. Smith, Frini Karayanidis

**Affiliations:** School of Science, College of Engineering, Science and Environment, University of Newcastle, Newcastle, New South Wales, Australia; Hunter Medical Research Institute, Newcastle, New South Wales, Australia; School of Engineering, College of Engineering, Science and Environment, University of Newcastle, Newcastle, New South Wales, Australia; Beckman Institute for Advanced Science and Technology, University of Illinois at Urbana-Champaign; Department of Psychology, University of Illinois Urbana-Champaign; School of Medicine and Public Health, College of Medicine, Health and Wellbeing, University of Newcastle, New South Wales, Australia; School of Psychology and Vision Sciences, University of Leicester, Leicester, UK; Alliance for Research in Exercise, Nutrition and Activity (ARENA), Allied Health and Human Performance (ARENA) Research Centre, College of Health, Adelaide University, South Australia, Australia

**Keywords:** Cerebral arterial elasticity, healthy ageing, cardiovascular risk factors, longitudinal effects

## Abstract

Ageing is often associated with a decline in cardiovascular and cerebrovascular health. This study examines the relationship between *cerebral* arterial elasticity and measures of cardiovascular health, as well as longitudinal changes in cerebral arterial elasticity over a 1.5 year interval. The pulse relaxation function (PReFx) is a measure of regional cerebral arterial elasticity derived using diffuse optical tomography (pulse-DOT). PReFx was measured over the anterior brain, including the frontal lobes and anterior sections of temporal and parietal regions that are especially vulnerable to vascular and cognitive ageing. We examined relationships between PReFx and measures of four cardiovascular risk factors (CVRF; i.e., hypertension, cholesterol, diabetes, obesity), as well as CVRF burden (i.e., number of CVRFs) in the highly active and cognitively healthy ACTIVate cohort (60-70 years). We replicated the well-established relationship between PReFx and both age and cardiorespiratory fitness, and examined associations between PReFx, measures of cardiovascular health and CVRF burden. Higher CVRF burden was linearly associated with lower cerebral arterial elasticity. The relationship between age and cerebral arterial elasticity was partially mediated by pulse pressure, an index of hypertension. PReFx declined significantly in as little as 1.5 years, and the effect did not vary with baseline level of any of the four CVRFs. We conclude that PReFx shows promise as a putative biomarker for monitoring cerebrovascular ageing in healthy older adults.

## 1. Introduction

Cardiovascular risk factors (CVRF) become more prevalent with increasing age and are associated with vascular changes that increase the risk of cardiovascular and cerebrovascular disease, such as heart disease and stroke. CVRF presence (e.g., hypertension, type 2 diabetes) is also associated with cognitive decline and increased dementia risk in older adults (Peters et al., 2020; Zimmerman et al., 2021). Characterising the relationship between subtle vascular changes that often precede the emergence of CVRFs and early signs of cognitive decline in mid-late life can inform and support development of targeted lifestyle and pharmaceutical intervention programs to prolong functional capacity and reduce risk of cognitive decline and dementia in old age.

A major contributor to age-related changes in cardiovascular health is arterial stiffening - the progressive thickening and hardening of arterial walls that results in less elastic arteries. Reduced arterial elasticity is one of the earliest manifestations of vascular ageing and increases exponentially with age (AlGhatrif et al., 2013; Najjar et al., 2005). Common measures of arterial elasticity, such as carotid-femoral pulse wave velocity (cf-PWV), focus on the large central vessels (e.g., aorta, carotid arteries) that are strongly impacted by changes in pulsatile blood flow (Badji et al., 2019). Arterial stiffening increases pulse pressure which accelerates atherosclerotic processes and increases endothelial damage and lipoprotein infiltration into the arterial wall (Kohn et al., 2015). Over the lifespan, this repeated mechanical pulse stress gradually alters arterial properties (e.g., increased collagen deposition, elastin fragmentation; O’Rourke and Hashimoto, 2007). These changes reduce the arterial wall’s ability to buffer pulsatile blood flow, leading to increased pulse pressure, elevated cardiac workload, and structural damage to sensitive end-organs such as the brain (Mitchell et al., 2011). The severity of change in arterial health varies considerably among mid-late life individuals, indicating that genetic, metabolic, behavioural and environmental factors affect rate of change across the lifespan (AlGhatrif et al., 2013).

There is evidence for bi-directional relationships between arterial stiffening and both cardiovascular and cerebrovascular health (e.g., Cecelja & Chowienczyk, 2013; Ikonomidis & Thymis, 2023; Jennings et al., 2020). Measures of high brachial blood pressure are associated with faster rate of arterial stiffening beyond that expected from ageing alone (Ohyama et al., 2016), even in prehypertensive individuals (AlGhatrif et al., 2013; see Wilson & Webb, 2020 for meta-analysis). Arterial stiffening predicts increased risk of incident hypertension at 8 years (Kaess et al., 2012), as well as high systolic blood pressure and hypertension risk at 4.5 years in normotensive middle-aged adults (Najjar et al., 2008). Across the lifespan, reduced arterial elasticity is associated with longitudinal increase in blood pressure and incident hypertension (Takase et al., 2011). Pulse pressure (i.e., the difference between systolic and diastolic brachial blood pressure) is considered a clinical indicator of arterial stiffening and cardio/cerebrovascular disease risk (Perdomo, 2019; Said et al., 2018). It is associated with cerebrovascular pathology (e.g., white matter lesions), cognitive decline, as well as increased risk of cardiovascular and cerebrovascular disease, and Alzheimer’s disease (Haider et al., 2003; Herzog et al., 2025).

Blood cholesterol level is also associated with increased arterial stiffness (Chen et al., 2021; Kim et al., 2010; Nabeel et al., 2021), especially in familial hypercholesterolemia (Kovács et al., 2022). The association between cholesterol level and cognitive ability varies across different measures of cholesterol with high density lipoprotein (HDL) showing the strongest relationship with general cognition and working memory performance (Georgina E. Crichton et al., 2014). The ratio of total cholesterol to HDL (Total/HDL) is a strong predictor of atherogenic risk (Bustamante Gallo et al., 2024), a stronger predictor of coronary disease than individual cholesterol measures in large cohort studies (e.g., (Castelli et al., 1986; Grover et al., 1994) and is recommended for use in clinical practice (Millán et al., 2009). This cholesterol ratio shows a non-linear relationship with cognition with high cholesterol ratio scores associated with poorer cognitive performance (Zhou et al., 2025).

Adiposity (Brunner et al., 2015) and type 2 diabetes (Elias et al., 2017) in late midlife are associated with higher rates of arterial stiffening over 4-5 years. Arterial elasticity predicted risk of type 2 diabetes at 4.5 year follow-up in older adults (Muhammad et al., 2017), and at 3.5 (Zheng et al., 2020) and 7-year follow-up (Cohen et al., 2022) in middle-aged adults. In fact, even pre-diabetic populations with impaired glucose tolerance show lower arterial elasticity (Prenner and Chirinos, 2015). Moreover, CVRFs often co-occur and the aggregation of multiple risk factors is associated with higher risk of cardiovascular disease (Cao et al., 2023) and 10-year mortality rate (Bazalar-Palacios et al., 2021). Scuteri et al. (2004) showed that metabolic syndrome, which represents the clustering of several CVRFs, is associated with greater vascular thickness and stiffening, independent of age. This strong evidence for complex, interactive relationships between arterial stiffening and CVRFs throughout adulthood has led to calls for monitoring arterial stiffening as an early indicator of cardiovascular risk, even in healthy individuals (Bazalar-Palacios et al., 2021).

Large artery stiffening is also associated with poorer performance on measures of global cognition as well as specific cognitive domains. Meta-analytic studies (e.g., Liu et al., 2021; Singer et al., 2014) report cross-sectional relationships between arterial stiffening and performance on cognitive screening tasks (e.g., Mini-Mental State Examination) as well as tasks tapping into cognitive domains sensitive to ageing (e.g., processing speed, executive functioning). A large meta-analysis of 29 cross-sectional and nine longitudinal studies (Alvarez-Bueno et al., 2020) showed that higher baseline arterial stiffening is associated with poorer performance on tests of global cognition as well as executive functioning and memory at baseline, and weaker but still significant longitudinal changes in these measures (Singer et al., 2014). Individual studies often reveal nuanced patterns of relationships between arterial elasticity and performance across cognitive domains. For instance, in middle-aged adults, low arterial elasticity was associated with poorer performance on the Trail-Making Task, but not other tasks of executive functioning, or tasks of visual processing or memory (Pase et al., 2016).

The relationship between large artery stiffening and cognition is also impacted by CVRF presence. For example, greater arterial stiffening combined with hypertension was associated with more severe cognitive impairment than either factor alone (Hajjar et al., 2016). Similarly, the association between arterial stiffening and global cognitive performance was stronger in individuals with multiple CVRF, and particularly in individuals with concurrent diabetes and hypertension (Elias et al., 2009). In LaPlume et al. (2022), adults aged 40-70 yrs with no risk factors performed cognitively at the level of people 10-20 years younger with multiple risk factors. Moreover, rate of cognitive decline was steeper in individuals with more CVRFs, particularly on executive function and processing speed tasks. This illustrates the protective effect of cardiovascular health on ageing and suggests that CVRF presence in young-mid adulthood may be linked to accelerated cognitive ageing.

These complex relationships between age, cardiovascular health, arterial stiffening and cognitive outcomes have been largely derived using carotid-femoral pulse wave velocity (cf-PWV), which primarily measures aortic elasticity. However, PWV measures cannot characterise subtle changes in the elasticity of cerebral arterial networks that perfuse brain regions sensitive to ageing and cerebrovascular health.

The diffuse optical tomography signal can be used to measure properties of the cerebral arterial pulse wave (pulse-DOT) that index cerebral arterial elasticity (Fabiani et al., 2014). Light sources on the scalp shine a spectrum of light that penetrates brain tissue (2-3cm). Some light is absorbed (depending on amount of haemoglobin present) and the remainder is reflected and picked up by detectors at some distance away on the scalp. Pulse-DOT measures are derived by analysing the pulse signal from intracranial arteries/arterioles which (like pulse oximetry) reflect absorption changes during a pulsation cycle. The pulse-DOT signal can be co-registered to the structural MRI to map regional variation in the elasticity in arteries and arterioles that perfuse different cortical regions.

The pulse relaxation function (PReFx) is a measure of arterial elasticity derived from the shape of the arterial pulse wave (Chiarelli et al., 2017). The shape is impacted by the temporal overlap between forward and reflected waves, representing how the artery recoils back to its original size after dilating during the systole. As arteries become stiffer, these waves overlap more, resulting in a smaller PReFx. PReFx is sensitive to ageing and ageing-associated changes in brain and cognition. In healthy middle and older age adults (55-87 years), a larger PReFx (indicative of more elastic arteries) was associated with higher cardiorespiratory fitness (eCRF, a composite estimate of VO₂ max; Jurca et al., 2005), as well as larger grey matter, white matter and total brain volume. In an adult lifespan cohort, higher PReFx in frontoparietal regions was associated with younger age and higher eCRF (Tan et al., 2017). In frontotemporal regions supplied by the middle cerebral artery (MCA), PReFx was associated with white matter lesions and fluid cognitive abilities (Tan et al., 2019).

Regional variability in arterial elasticity is also associated with distinct cognitive processes. Tan et al. (2017) found that working memory performance was positively correlated with PReFx in frontoparietal regions, but not across the whole brain or in task-irrelevant areas (e.g., visual cortex). Other arterial elasticity measures derived from the pulse-DOT signal show similar findings. For example, longer pulse transit time (PTT, the time for the pulse wave to travel from the heart to the point of measurement) in the left MCA which perfuses Broca’s area, was associated with better performance on verbal fluency but not working memory tasks (Fabiani et al., 2014). Conversely, longer PTT in areas including the dorsolateral prefrontal cortex (dlPFC), which are perfused by the superior portion of the precentral artery bilaterally, was associated with better performance on working memory but not verbal fluency tasks. In a cross-sectional lifespan sample, a decline in PreFx emerged almost 10 years earlier than in white matter lesion volume (Bowie et al., 2024). Moreover, the effects of age on fluid cognitive abilities (but not crystallised abilities) were mediated by variability in PReFx as well as white matter health. These findings are consistent with hierarchical cascade models (e.g., Kong et al., 2020) which posit that subtle progressive declines in arterial elasticity can contribute to structural and functional brain changes in the areas they perfuse, which, in turn, lead to decline in the specific cognitive functions that these areas support.

### 1.1 Present Study

This study seeks to characterise the relationship between cerebral arterial elasticity and indicators of cardiovascular health in the ACTIVate cohort, a large sample of cognitively healthy, community-dwelling 60-70yr old Australians (Smith et al., 2022). Cerebral arterial elasticity was measured using PReFx over the anterior brain encompassing the frontal lobes and anterior sections of the temporal and parietal lobes that are particularly vulnerable to both ageing and vascular changes.

In the baseline cross-sectional cohort, we first investigated whether cerebral arterial elasticity over the anterior brain varies with increasing burden of four prevalent CVRFs (i.e., hypertension, cholesterol, diabetes, obesity) that have potentially synergistic effects on vascular and cognitive health (e.g., LaPlume et al., 2022). We then explored whether clinical indicators of these four CVRFs mediate the well-established relationship between age and cerebral arterial elasticity (Fabiani et al., 2014; Tan et al., 2017).

In addition, for the first time, we investigated whether cerebral arterial health changes over a 1.5 year follow-up period in a subsample of the ACTIVate cohort. Large artery stiffening increases with age with steeper declines emerging over longer timeframes (e.g., AlGhatrif et al., 2013; Najjar et al., 2008). The rate of aortic stiffening is accelerated by the presence of CVRFs even over relatively short periods (e.g., <6 years, Benetos et al., 2002; Crichton et al., 2014). Longitudinal studies show evidence of a small, but measurable, annual rate of decline in brain structure (e.g., Fujita et al., 2023) as well as global cognitive ability (e.g., Zaninotto et al., 2018) in older adults. We argue that a decline in cerebral arterial elasticity over this relatively brief temporal window, especially in community-dwelling, healthy older adults, would provide strong preliminary evidence for PReFx as a potential biomarker of cerebral arterial health. Finally, we also explored whether PReFx change over this 1.5 year period varies with level of CVRF burden or level of clinical indicators of CVRFs.

## 2. Methods

### 2.1. Participants

The Newcastle arm of the ACTIVate Study included community-dwelling older adults (*N* = 200, 60-70 years, *M ± SD:* 65.5 ± 3.2 yrs, 57% female) were recruited through the Hunter Medical Research Institute’s (HMRI) Volunteer Registry, Facebook advertisements and word of mouth (for full protocol see Smith et al., 2022). Ethics approval was obtained from the University of South Australia (202639) and registered with the University of Newcastle Human Research Ethics Committees. All procedures were conducted in accordance with the Declaration of Helsinki and participants gave written informed consent.

At Baseline (Phase 1), 173 had complete pulse-DOT data sets. Of these, eight did not pass quality checks, and two were classified as outliers and removed (see below). Therefore, cross-sectional data are reported from 163 participants (65.4 ± 3.1 yrs, 59% female).

All 180 participants who returned for 1.5 yr follow-up testing (Phase 2) were also invited to complete the optical imaging component by extending the testing session or returning at another date^1^. Ninety two participants attended the second session. Of these, due to technical issues or poor data quality, 64 participants (70%) had complete pulse-DOT datasets at both phases (Phase 1: 65.2 ± 3.1 yrs, Phase 2: 66.8 ± 3.1 yrs; 58% female) with an average 19 ± 1.6 month follow-up period. There were no baseline differences in demographic, cognitive and physiological variables between this subgroup and the remainder of the cohort who only had Phase 1 optical data (smallest *p* = .273).

#### 2.1.1. Inclusion/exclusion criteria

At baseline, participants were included if they were aged 60-70 years at screening and fluent English-speakers. Participants were excluded if they reported a current clinical diagnosis of dementia, scored in the dementia range on the Montreal Cognitive Assessment-Blind (MoCA-B delivered over the phone, <13/22; Wittich et al., 2010), were colour blind or had an uncorrected visual condition, or reported diagnosis of a major neurological (including stroke or transient ischaemic attack) or psychiatric condition, a known intellectual or major physical disability, a previous head trauma resulting in loss of consciousness of more than five minutes, a current or previous alcohol or substance dependence, recreational drug use in the last three months or cancer treatment affecting cognitive performance (e.g. chemotherapy) in the last five years. Participants did not complete brain imaging, if they did not meet MRI inclusion criteria (i.e., had a pacemaker, metal implants or fragments, or a history of claustrophobia).

### 2.2. Measures

#### 2.2.1. Cognitive functioning

was estimated using the total score on Addenbrooke’s Cognitive Examination III (ACE-III; Hsieh et al., 2013) which assesses cognitive function across five domains: memory, attention, language, fluency, and visuospatial ability.

#### 2.2.2. Physical activity level

was measured using a triaxial accelerometer (Axivity AX3) sampling at 100 Hz over seven consecutive days on the non-dominant wrist. Daytime behaviours were classified as moderate to vigorous physical activity (MVPA; > 93 milligravity units (mg)), light intensity physical activity (LPA; >48mg) or sedentary (<48mg) (Mellow et al., 2024). Moderate-to-vigorous physical activity (MVPA) was quantified as the average number of minutes of MVPA/day. Data were pre-processed using Open Movement GUI software (OmGUI) and analysed using a custom MATLAB toolbox (COBRA; MATLAB R2018b; see Mellow et al., 2022, 2024 for analysis protocol).

#### 2.2.3. Cardiovascular measures

were measured using protocols described in Smith et al. (2022).

Systolic and diastolic blood pressure (mmHg) and heart rate (beats per minute) were measured as the average across three consecutive brachial blood pressure measurements with ∼3min break. Pulse pressure was calculated as the difference between systolic and diastolic blood pressure.

Level of total cholesterol, high-density lipoprotein cholesterol (HDL), low-density lipoprotein cholesterol (LDL), glucose and triglycerides (all expressed in mmol/L) were measured in fasting venous blood samples. Samples were obtained through venipuncture, centrifuged at 4000 rpm for 10 minutes and stored at −80 degrees Celsius. Analyses were conducted using a KONELAB 20XTi auto-analyzer and Thermo Fisher reagents. Body mass index (BMI, kg/m²) was calculated using the average of two measurements of height and weight.

#### 2.2.4. Estimated cardiorespiratory fitness (eCRF)

was calculated by linearly combining weighted variables for sex, age, BMI, resting heart rate, and self-reported physical activity (Jurca et al., 2005). eCRF is highly predictive of VO₂Max and has been validated in older adults (Mailey et al., 2010). eCRF uses a constant to correct for systematic differences in lung capacity between males and females when estimating VO₂Max. When including eCRF in analyses with other variables, we removed the correction of sex, as the sex-corrected score does not reflect real variations in vascular health (Fabiani et al., 2014).

#### 2.2.5. CVRF presence and burden level

were operationalised using four metabolic indicators of CVRFs: blood pressure, blood cholesterol, blood glucose and BMI. These were selected because hypertension, hypercholesteremia, diabetes and obesity are common amongst older adults in Western cultures and strongly associated with increased risk of cardiovascular disease, cognitive decline and dementia (Cao et al., 2023; Farnsworth Von Cederwald et al., 2022).

CVRF burden was defined as the number of CVRFs present at Phase 1. Each of four CVRFs was characterised as present or absent using Australian guideline thresholds for each measure (Nelson et al., 2024) and participant self-report of a diagnosis. Very few people were classified as having all four CVRFs, resulting in four levels of CVRF burden (0, 1, 2, 3+).

Hypertension was classified as present if the participant had systolic blood pressure (sBP) was above 140 mmHg, diastolic blood pressure (dBP) above 90 mmHg, reported a clinical diagnosis of hypertension or was taking one or more antihypertensive medication^2^. High cholesterol was classified as present if the participant had fasting total blood cholesterol over 6.5 mmol/L (i.e., 251 mg/dL), high-density lipoprotein below 1.5 mmol/L (i.e., 58 mg/dL), reported having received a clinical diagnosis of high blood cholesterol or was taking one or more cholesterol lowering medication. Diabetes was classified present if the participant had a fasted blood glucose level above 7.0 mmol/L (i.e., 126 mg/dL), reported having received a clinical diagnosis of diabetes or was taking one or more blood glucose lowering medication. Indication of each medication classification was confirmed by a pharmacist. Obesity was classified as present if BMI exceeded 30 kg/m², which is more commonly reported than waist-hip ratio (WHR)^3^.

To explore the potential mediating effects of these CVRF factors on the relationship between age and cerebral arterial elasticity, we defined a single quantitative measure for each CVRF. For hypertension, we used pulse pressure (systolic-diastolic brachial blood pressure). Pulse pressure is a stronger predictor of cardiovascular events than systolic or diastolic blood pressure alone, especially in older adults (Benetos et al., 1997; Mancusi et al., 2018), and a clinical indicator of arterial stiffening and vascular disease risk (Perdomo, 2019; Said et al., 2018). For cholesterol, we used cholesterol ratio (Total/HDL) which represents a higher unfavourable proportion of unhealthy to protective cholesterol. Cholesterol ratio is a strong predictor of risk of atherosclerosis and coronary disease (Bustamante Gallo et al., 2024; Castelli et al., 1986; Grover et al., 1994), recommended for clinical practice and is used in cardiovascular risk calculators (Millán et al., 2009). Blood glucose level was used a measure of diabetes risk, and BMI as a measure of obesity.

#### 2.2.6. Diffuse Optical Tomography

Optical data were recorded with a multi-channel frequency-domain oximeter (Imagent; ISS Inc., Champaign, IL USA) using 4 detectors and 16 time-multiplexed light emitting diodes generating light at 830 and 690 nm (max amplitude 10mW, mean amplitude after multiplexing: 1 mW) modulated at 110 MHz) resulting in 128 channels (i.e., source-detector pairings). Sources and detectors were held against the participant’s scalp using a custom-built, soft foam, adjustable helmet. The light was transmitted to the scalp via optical fibres (diameter = 400 µm; one fibre per emitter) with paired fibres carrying each wavelength to each location. Detector fibre bundles (diameter = 3 mm) collected light from the scalp and were connected to photomultiplier tubes (PMTs) fed with a current modulated at 110.003125 MHz, generating a 3.125 kHz cross-correlation frequency. A Fast Fourier Transform of the PMT output data was used to calculate direct current (DC) intensity, alternating current (AC) intensity, and relative phase. Optical parameters were sampled at 39.0625 Hz (25.6 ms per sampling point).

Data were acquired from three optical montages recorded consecutively over left, middle and right anterior brain in that order, with each montage covering areas of the frontal cortex, as well as anterior temporal and parietal cortices. Across montages, this resulted in a total of 384 channels (i.e., source-detector pairs). For each montage, four blocks of data were recorded (3 min/block): a resting block, two breath-holding task blocks and a second resting block. The breath-holding data are not reported here.

### Electrocardiogram (ECG)

data were used to synchronize the pulse-DOT data to the arterial pulse wave. ECG was recorded from electrodes attached to the left and right wrist with a sampling rate of 1000 Hz (AD Instruments Power Series 25). The optical pulse data were time-locked to the R-wave of the ECG to ensure the same pulse was examined regardless of spatial location of optical signal. R-peak detection was implemented using an algorithm running on MATLAB R2021b (MathWorks) that used a band-pass filter of 1-50 Hz, searched for peaks exceeding a dynamic voltage threshold, and discarded points that occurred outside the normal range of interbeat intervals. Accurate identification of each R-peak was ensured by visual inspection, and false detections (e.g., a large T wave inadvertently identified as an R wave) were manually corrected or eliminated.

### Structural magnetic resonance imaging (MRI)

data were used to align the optical data to brain anatomy (sMRI). T1-weighted structural scans were collected on a Siemens Prisma 3T scanner using a 64-channel head and neck coil (MPRAGE sequence; voxel size = 1×1×1 mm³; TE = 2.91 ms; TR = 2300 ms; TI = 900 ms; flip angle = 9°). These scans were used for co-registration of optical imaging data.

### Digitisation was used for co-registration with structural MRI

The spatial location of source and detector positions on the optical helmet as well as the nasion and preauricular reference points were digitised individually for each participant using a recording stylus and three head-mounted receivers, which allowed for small movements of the head between measurements (Polhemus 3Space: FASTRAK 3D digitizer, Polhemus, Colchester, VT). The Polhemus digitisation points were co-registered to the individual’s structural MRI using in-house software (Optimized Co-registration Package, OCP; Chiarelli et al., 2015). The helmet was worn continuously during the digitisation and recording session and participants were instructed to avoid large head movements.

### Measurement of pulse parameters

We report analyses from AC intensity values at 830 nm only (Fabiani et al., 2014), resulting in 192 channels across the three montages (16 sources by 4 detectors) covering fronto-temporo-parietal regions. Data were analysed separately for each resting state block to examine reliability across blocks and then averaged for final analyses (see below).

The optical data were normalised (by dividing by their mean value) and band-pass filtered between 0-10Hz. *Opt-3d* software (Gratton, 2000) was used to combine channels whose mean diffusion paths overlapped for a given brain voxel. For each channel, each voxel is assigned a weight related to the estimated sensitivity of the measures to phenomena (absorption and scattering) occurring in brain regions underlying that voxel. This sensitivity was estimated based on a perturbation model (Feng et al., 1995), such that a single waveform was created for each voxel from a weighted sum of all the channel waveforms whose diffusion path was included in that voxel. To reduce sensitivity to superficial artefacts and increase the signal-to-noise ratio (SNR), only channels with source-detector distances between 20-60 mm were included permitting phenomena approximately 1-3 cm in depth to be imaged. The voxel data were surface projected (2D model) to the superior axial and left/right sagittal surfaces. Waveforms were time-locked to the peak of the ECG R wave to ensure the same pulse cycle was measured in all locations.

Three regions of interest (ROI) were defined using the Automated Anatomical Labelling Atlas 3 (AAL3, Rolls et al., 2020) to comprise voxels covering the frontal cortex and portions of the temporal and parietal cortices that are perfused by branches of the left and right middle cerebral arteries (l-MCA, r-MCA) and anterior cerebral artery (ACA, **Error! Reference source not found.**) at a max depth of 2-3 cm.

The Pulse Relaxation Function (PReFx) was derived from each ROI using Opt-3d software (Gratton, 2000). PReFx is a measure of the shape of the pulse wave and is calculated by determining the area beneath the pulse waveform between the peak systole and peak diastole after subtracting a triangular area that represents “linear” relaxation. This value is normalised by setting the peak systole to 1 and peak diastole to 0 and dividing them by the duration of the systole-diastole interval. This ensures that PReFx is independent of pulse amplitude and heart rate.

Individual trials (i.e., pulse cycles) for each ROI and each block were discarded if: (1) no systolic or diastolic peaks were identified within the epoch; (2) PReFx value was greater than 1.0 or less than −0.2; (3) peak amplitude was less than 0.001 dB or greater than 3 dB; (4) the interbeat interval (IBI) estimate based on the optical data differed from the IBI obtained from the ECG by more than 76.8 ms (3 sampling points); or (5) the IBI estimate was smaller than 500 ms (>120 beats per minute) or larger than 1500 ms (<40 beats per minute). For each ROI, at least 10% of voxels were required to have a minimum of 10 valid trials. ROIs that did not meet this criterion were excluded from further analysis. For remaining trials, PReFx was averaged across all voxels in each ROI for each block of the two rest blocks (PReFx at l-MCA, ACA and r-MCA). PReFx values were also averaged across all three ROIs (Total PFC).

At Phase 1, eight participants had one or more ROIs that did not pass these checks and were removed. For each participant and each ROI, outlier PReFx values were identified by calculating the difference in PReFx across the two rest blocks. If this difference was ±2.96 *SD* from the sample mean, we examined the shortest channels within that ROI. If a clear pulse could not be identified, this ROI for this participant was considered an outlier and removed. This resulted in removal of all ROI data for two participants. Thus, at Phase 1, PReFx data were available for analyses from 163 participants.

### 2.3. Procedure

Phase 1 (baseline) data were collected over three testing sessions over roughly a two-week period. Phase 2 data were collected approximately 1.5 years after baseline assessment with some participants returning for a second optical session. MRI and bloods were not collected at Phase 2.

### 2.4. Data Analyses

The data were analysed and visualised in R - version 4.3.1 (R Core Team, 2025) and SPSS v30. PReFx reliability was analysed using Pearson correlations (2-tailed) to confirm PReFx test-retest reliability across the two resting state blocks at each ROI and reliability across the three ROIs after averaging across the resting-state blocks within each ROI. Laterality differences in PReFx across the three regions of interest (ROIs: l-MCA, ACA, r-MCA) was examined using a repeated measures GLM with post-hoc pairwise comparisons.

#### 2.4.1. Cross-sectional data

Cohort profile at baseline was characterised using demographic measures and indicators of cardiovascular health compared against Australian norms.

We used Pearson correlations (a=0.05, 2-tailed, bootstrapped 1000 iterations) with false discovery rate (FDR) correction (Benjamini and Hochberg, 1995; Yekutieli and Benjamini, 1999) to examine relationships between demographic, cognitive, cardiovascular health measures and PReFx values at each ROI (l-MCA, ACA and r-MCA).

We examined whether PReFx varied with CVRF Burden using a 4 CVRF Burden (0, 1, 2, 3+) x 3 ROI mixed-design GLM with polynomial group trends.

Finally, we explored whether the level of each of the four CVRFs (i.e., pulse pressure, cholesterol ratio, glucose, BMI) mediated the relationship between age and PReFx, using mediation models (PROCESS macro for R, Model 4; Hayes, 2022). Bootstrapping (10,000 iterations) was used to generate 95% confidence intervals for all indirect effects.

#### 2.4.2. Longitudinal data

We examined whether PReFx declined over time using a 2 Phase (Baseline, Follow-up) x 3 ROI (l-MCA, ACA, r-MCA) repeated measures ANOVA. Post-hoc comparisons were used to examine PReFx decline at each ROI using paired t-tests (2-tailed, bootstrapped 1000 iterations).

As the decline in PReFx was only significant at ACA and r-MCA, we ran further exploratory analyses to characterise whether the decline in PReFx varied with cardiovascular burden at baseline. We used a 4 CVRF-Burden (0, 1, 2, 3+) x 2 ROI (ACA, r-MCA) mixed design ANOVA on PReFx change scores (Baseline - Follow-up).

Finally, we explored whether age and baseline level of key cardiovascular risk factors (i.e., pulse pressure, cholesterol ratio, BMI and glucose) were associated with the level of PReFx change at either ACA or r-MCA (Pearson pairwise correlations, 2-tailed, bootstrapped 10,000 iterations with FDR correction).

## 3. Results

### 3.1. Cross-sectional analyses: Phase 1 (Baseline)

#### 3.1.1. Cohort profile

**Error! Reference source not found.** shows demographic, cognitive and cardiovascular health variables at baseline. The sample was highly educated, with 59% participants having completed post-secondary education. Total ACE-III score was high with only one participating below the dementia cut-off (<82/100; Hsieh et al., 2013)^4^, as expected given cognitive screening using the MoCA-B (Wittich et al., 2010).

Using Australian guidelines (Nelson et al., 2024), on average, the cohort scored within or just above the healthy cut-off values on measures for all four CVRFs (Table 1). On average, the cohort was highly active, averaging almost 1.5 hrs of moderate-to-vigorous physical activity (MVPA) per day. However, there was very substantial variability, with MVPA ranging from 3.4 min to over 3 hours per day.

**Table 1.** Key cohort measures at Phase 1 (Baseline).

| **Variable** | **N** | **M ±SD** | **Range** | **Men** | **Women** | **p** |
| --- | --- | --- | --- | --- | --- | --- |
| *Age (yrs)* | 163 | 65.4 ±3.1 | 60.1-71.2 | 65.9 | 65.0 | .080 |
| *Education (yrs)* | 161 | 16.8 ±3.1 | 7-24 | 16.9 | 16.6 | .609 |
| *Addenbrooke's Cognitive Examination III (ACE-III)* | 163 | 94.2 ±3.8 | 75-100 | 93.5 | 94.7 | .058 |
| *Systolic Blood Pressure (sBP; >140 mmHg\*)* | 159 | 141.5 ±17.1 | 106.3- 188.0 | 146.7 | 137.9 | .001 |
| *Diastolic Blood Pressure (dBP; >90 mmHg)* | 159 | 77.8 ±8.7 | 57.7-102.3 | 79.3 | 76.7 | .064 |
| *Pulse Pressure (PP; >60 mmHg)* | 159 | 63.7 ±13.2 | 35.7-103.7 | 67.4 | 61.2 | .004 |
| *Heart Rate (healthy range 60-100 beats/min)* | 159 | 64.9 ±10.8 | 42.7-107.7 | 62.3 | 66.7 | .011 |
| *Total-Cholesterol (Chol; >6.5 mmol/L)* | 158 | 5.4 ±1.1 | 2.3-7.9 | 5.1 | 5.6 | <.004 |
| *High Density Lipo-protein (HDL; <1.5 mmol/L)* | 158 | 1.7 ±0.5 | 0.5-3.1 | 1.5 | 1.8 | <.001 |
| *Cholesterol Ratio (Chol/HDL; <6.0 mmol/L)* | 158 | 3.4 ±1.3 | 1.6-11.4 | 3.6 | 3.3 | .212 |
| *Low Density Lipo-protein (LDL; >1.8 mmol/L)* | 158 | 3.2 ±1.0 | 0.8-6.2 | 3.0 | 3.3 | .154 |
| *Triglycerides (> 2.0 mmol/L)* | 158 | 1.2 ±0.7 | 0.2-6.0 | 1.2 | 1.2 | .760 |
| *Glucose (>7.0 mmol/L)* | 158 | 4.9 ±0.7 | 3.7-8.1 | 5.1 | 4.8 | .003 |
| *Body Mass Index (BMI; >30)* | 161 | 27.2 ±5.1 | 17.1-43.8 | 27.7 | 26.9 | .296 |
| *Mod-vigorous Physical Activity (MVPA min/day)* | 152 | 88.8 ±48.6 | 3.4-229.7 | 106.1 | 75.5 | <.001 |
| *Estimated cardiorespiratory fitness (eCRF; MET)* | 156 | 8.0 ±2.4 | 2.5-12.5 | 9.6 | 6.8 | <.001 |

*Note: Clinical cut-offs are based on Australian guidelines (Nelson et al. 2024). eCRF is reported with sex adjustment here to represent MET score. The sex effect is not significant in unadjusted scores that are used in subsequent analyses.*

Overall, 57% of the cohort was classified as having hypertension, 83% as having high cholesterol, but only 5% and 27% had evidence of diabetes or obesity, respectively (Table 2). Only 8% of the cohort had no CVRFs, whereas 19% had three or more CVRF, and roughly equivalent percentage having one or two CVRFs. Females were less likely to have hypertension than males (*X*^2^(1, *N* = 155) = 4.96, *p* = .003) but there were no other sex differences in either CVRF type or CVRF Burden. There was no age difference between CVRF Burden groups.

**Table 2.**
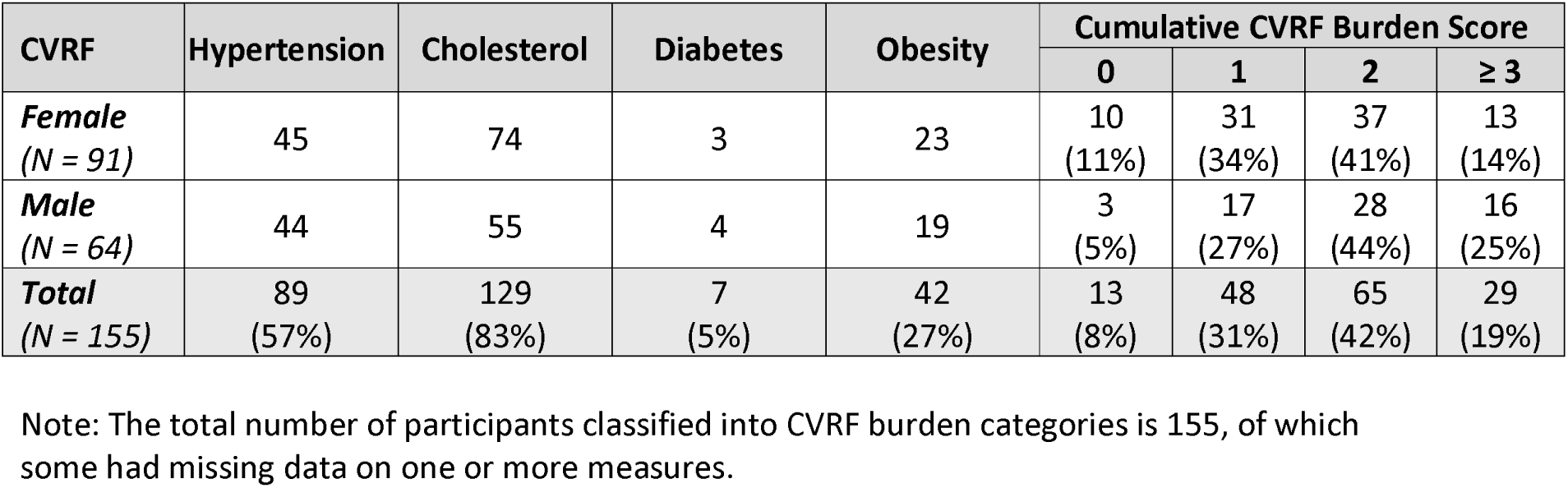
Distribution of CVRF presence and burden at Baseline.

| CVRF | Hypertension | Cholesterol | Diabetes | Obesity | Cumulative CVRF Burden Score |  |  |  |
| --- | --- | --- | --- | --- | --- | --- | --- | --- |
|  |  |  |  |  | 0 | 1 | 2 | ≥ 3 |
| **Female**<br>(N = 91) | 45 | 74 | 3 | 23 | 10<br>(11%) | 31<br>(34%) | 37<br>(41%) | 13<br>(14%) |
| **Male**<br>(N = 64) | 44 | 55 | 4 | 19 | 3<br>(5%) | 17<br>(27%) | 28<br>(44%) | 16<br>(25%) |
| **Total**<br>(N = 155) | 89<br>(57%) | 129<br>(83%) | 7<br>(5%) | 42<br>(27%) | 13<br>(8%) | 48<br>(31%) | 65<br>(42%) | 29<br>(19%) |

Note: The total number of participants classified into CVRF burden categories is 155, of which some had missing data on one or more measures.

#### 3.1.2. Relationships between age, cognition and cardiovascular health

Table 3A presents Pearson’s correlations between age, cognition, and cardiovascular health variables.

**Table 3.**
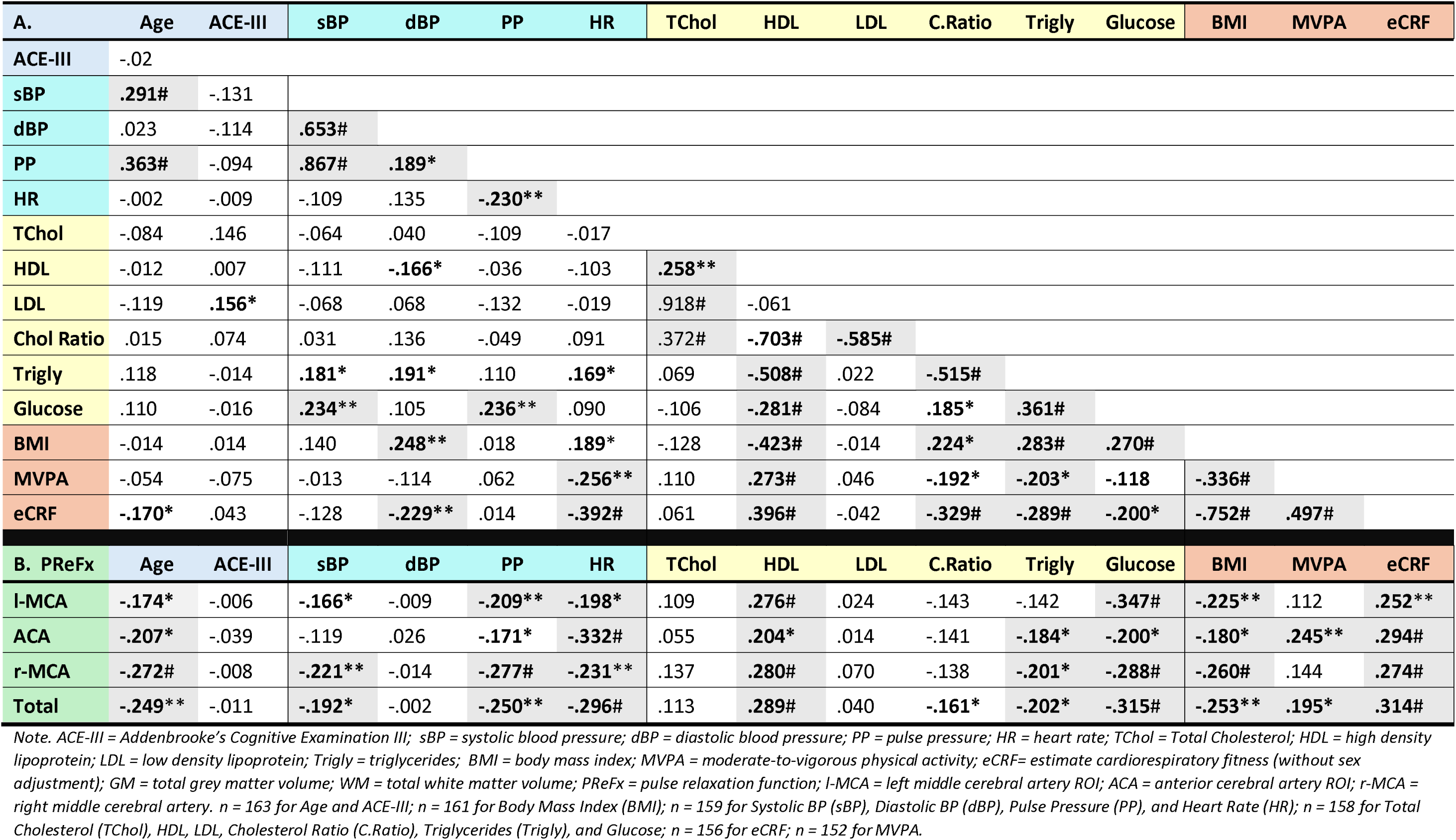
A. Correlations between key demographic, cognitive and cardiovascular health measures. **B.** Correlations between PReFx measures for each ROI and key demographic, cognitive and cardiovascular health measures. r-values from Pearson correlations (2-tailed). Bold indicates significant effect (* p ≤ .05; ** p ≤ .01; # p ≤ .001). Grey shading indicates that relationships that survived FDR correction.

| A. | Age | ACE-III | sBP | dBp | PP | HR | TChol | HDL | LDL | C.Ratio | Trigly | Glucose | BMI | MVPA | eCRF |
| --- | --- | --- | --- | --- | --- | --- | --- | --- | --- | --- | --- | --- | --- | --- | --- |
| ACE-III | -.02 |  |  |  |  |  |  |  |  |  |  |  |  |  |  |
| sBP | **.291#** | -.131 |  |  |  |  |  |  |  |  |  |  |  |  |  |
| dBp | .023 | -.114 | **.653#** |  |  |  |  |  |  |  |  |  |  |  |  |
| PP | **.363#** | -.094 | **.867#** | **.189\*** |  |  |  |  |  |  |  |  |  |  |  |
| HR | -.002 | -.009 | -.109 | .135 | **-.230\*\*** |  |  |  |  |  |  |  |  |  |  |
| TChol | -.084 | .146 | -.064 | .040 | -.109 | -.017 |  |  |  |  |  |  |  |  |  |
| HDL | -.012 | .007 | -.111 | **-.166\*** | -.036 | -.103 | **.258\*\*** |  |  |  |  |  |  |  |  |
| LDL | -.119 | **.156\*** | -.068 | .068 | -.132 | -.019 | .918# | -.061 |  |  |  |  |  |  |  |
| Chol Ratio | .015 | .074 | .031 | .136 | -.049 | .091 | .372# | **-.703#** | **-.585#** |  |  |  |  |  |  |
| Trigly | .118 | -.014 | **.181\*** | **.191\*** | .110 | **.169\*** | .069 | **-.508#** | .022 | **-.515#** |  |  |  |  |  |
| Glucose | .110 | -.016 | **.234\*\*** | .105 | **.236\*\*** | .090 | -.106 | **-.281#** | -.084 | **.185\*** | **.361#** |  |  |  |  |
| BMI | -.014 | .014 | .140 | **.248\*\*** | .018 | **.189\*** | -.128 | **-.423#** | -.014 | **.224\*** | **.283#** | **.270#** |  |  |  |
| MVPA | -.054 | -.075 | -.013 | -.114 | .062 | **-.256\*\*** | .110 | **.273#** | .046 | **-.192\*** | **-.203\*** | **-.118** | **-.336#** |  |  |
| eCRF | **-.170\*** | .043 | -.128 | **-.229\*\*** | .014 | **-.392#** | .061 | **.396#** | -.042 | **-.329#** | **-.289#** | **-.200\*** | **-.752#** | **.497#** |  |
| B. PReFx | Age | ACE-III | sBP | dBp | PP | HR | TChol | HDL | LDL | C.Ratio | Trigly | Glucose | BMI | MVPA | eCRF |
| I-MCA | **-.174\*** | -.006 | **-.166\*** | -.009 | **-.209\*\*** | **-.198\*** | .109 | **.276#** | .024 | -.143 | -.142 | **-.347#** | **-.225\*\*** | .112 | **.252\*\*** |
| ACA | **-.207\*** | -.039 | -.119 | .026 | **-.171\*** | **-.332#** | .055 | **.204\*** | .014 | -.141 | **-.184\*** | **-.200\*** | **-.180\*** | **.245\*\*** | **.294#** |
| r-MCA | **-.272#** | -.008 | **-.221\*\*** | -.014 | **-.277#** | **-.231\*\*** | .137 | **.280#** | .070 | -.138 | **-.201\*** | **-.288#** | **-.260#** | .144 | **.274#** |
| Total | **-.249\*\*** | -.011 | **-.192\*** | -.002 | **-.250\*\*** | **-.296#** | .113 | **.289#** | .040 | **-.161\*** | **-.202\*** | **-.315#** | **-.253\*\*** | **.195\*** | **.314#** |

Note. ACE-III = Addenbrooke's Cognitive Examination III; sBP = systolic blood pressure; dBp = diastolic blood pressure; PP = pulse pressure; HR = heart rate; TChol = Total Cholesterol; HDL = high density lipoprotein; LDL = low density lipoprotein; Trigly = triglycerides; BMI = body mass index; MVPA = moderate-to-vigorous physical activity; eCRF = estimate cardiorespiratory fitness (without sex adjustment); GM = total grey matter volume; WM = total white matter volume; PReFx = pulse relaxation function; I-MCA = left middle cerebral artery ROI; ACA = anterior cerebral artery ROI; r-MCA = right middle cerebral artery. $n = 163$ for Age and ACE-III; $n = 161$ for Body Mass Index (BMI); $n = 159$ for Systolic BP (sBP), Diastolic BP (dBp), Pulse Pressure (PP), and Heart Rate (HR); $n = 158$ for Total Cholesterol (TChol), HDL, LDL, Cholesterol Ratio (C.Ratio), Triglycerides (Trigly), and Glucose; $n = 156$ for eCRF; $n = 152$ for MVPA.

Even within this 10-year restricted age range, increasing age was associated with significantly higher systolic blood pressure and pulse pressure, but not other measures of cardiovascular health. ACE-III scores showed a small *positive* correlation with LDL cholesterol level, but this did not survive FDR correction.

Correlations between blood pressure, heart rate, cholesterol, triglycerides, glucose and obesity measures of cardiovascular health showed expected patterns, with groupings most strongly within measures tapping into similar mechanisms.

Consistent with eCRF being a reliable estimate of cardiorespiratory fitness, higher eCRF was associated with significantly lower dBP, heart rate, cholesterol ratio, triglycerides and glucose, BMI and higher HDL cholesterol and MVPA. MVPA showed a similar pattern but overall weaker correlations.

#### 3.1.3. PReFx reliability and regional differences

PReFx showed strong test-retest reliability between first and second recording blocks for all ROIs: l-MCA *(r*(161) = .857, *p* ≤ .001), ACA (*r*(161) = .887, *p* < .001), and r-MCA (*r*(161) = .863, *p* ≤ .001). For all subsequent analyses, we report data averaged across the two resting state blocks.

Figure 2 shows PReFx scores at l-MCA, ACA and r-MCA, as well as the average across the three ROIs (Total-PFC). PReFx was moderately to strongly positively correlated across the three ROIs (*r*(161) = .590 - .698, all *p* ≤ .001). Mean PReFx differed significantly between the three ROIs (*F*(2,324) = 10.03, *p* ≤.001). Post-hoc comparisons indicated that PReFx was significantly lower in the l-MCA compared to both the ACA and r-MCA (*F*(1,162) = 16.46, *p* ≤ .001; *F*(1,162) = 10.51, *p* ≤ .001, respectively) but did not differ between the ACA and r-MCA (*F*(1,162) = 2.39, *p* = .124). These findings indicate that PReFx measures were highly reliable across ROIs but still showed regional variability.

**Figure 1.**
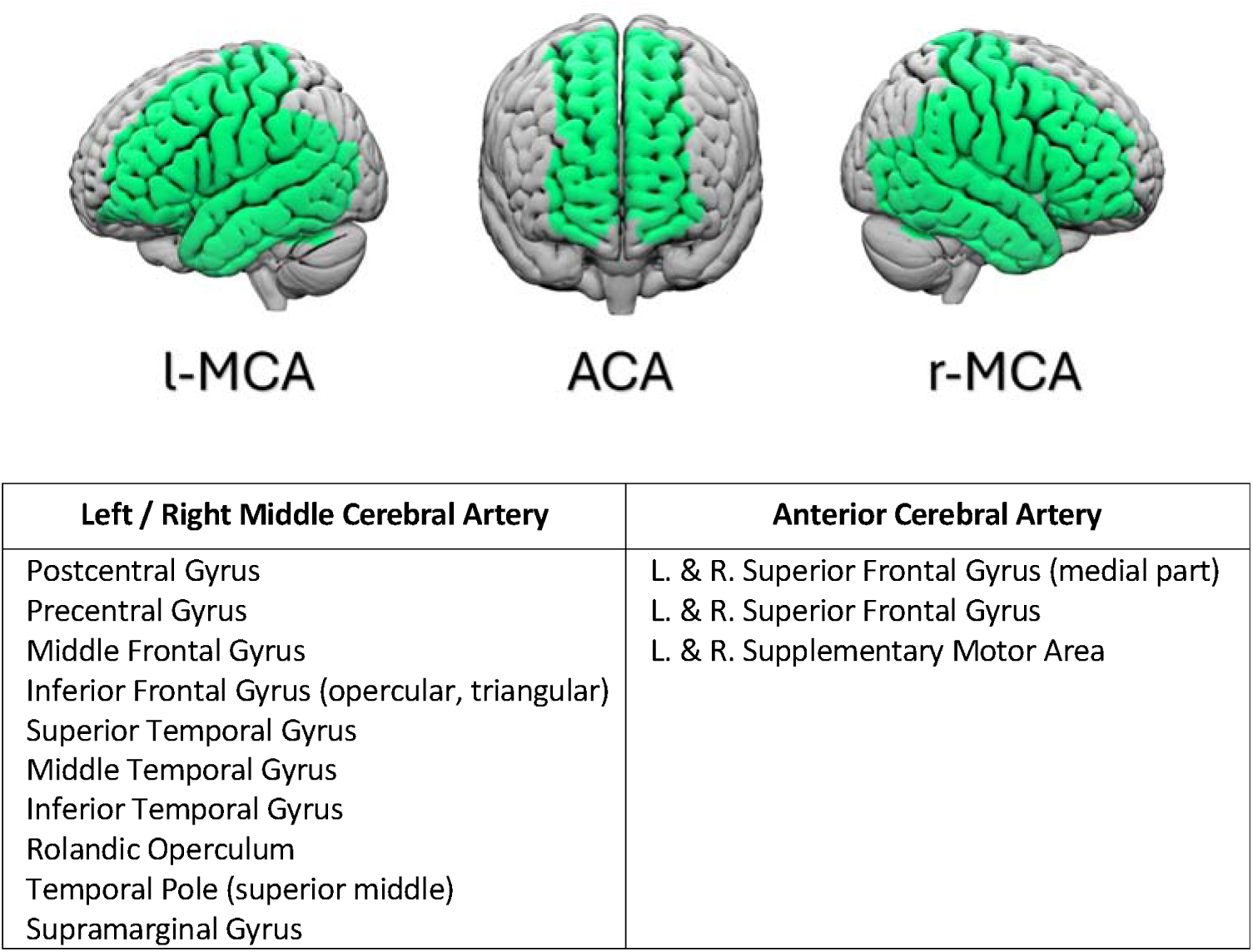
Pulse-DOT recording montages. Green areas represent that areas from anterior brain areas perfused by the left and right Middle Cerebral Artery (i.e. l-MCA, r-MCA) and midline (i.e. ACA) from which optical signal was acquired. Visualised in ‘Surfice’ (Rorden, 2025).

**Figure 2.**
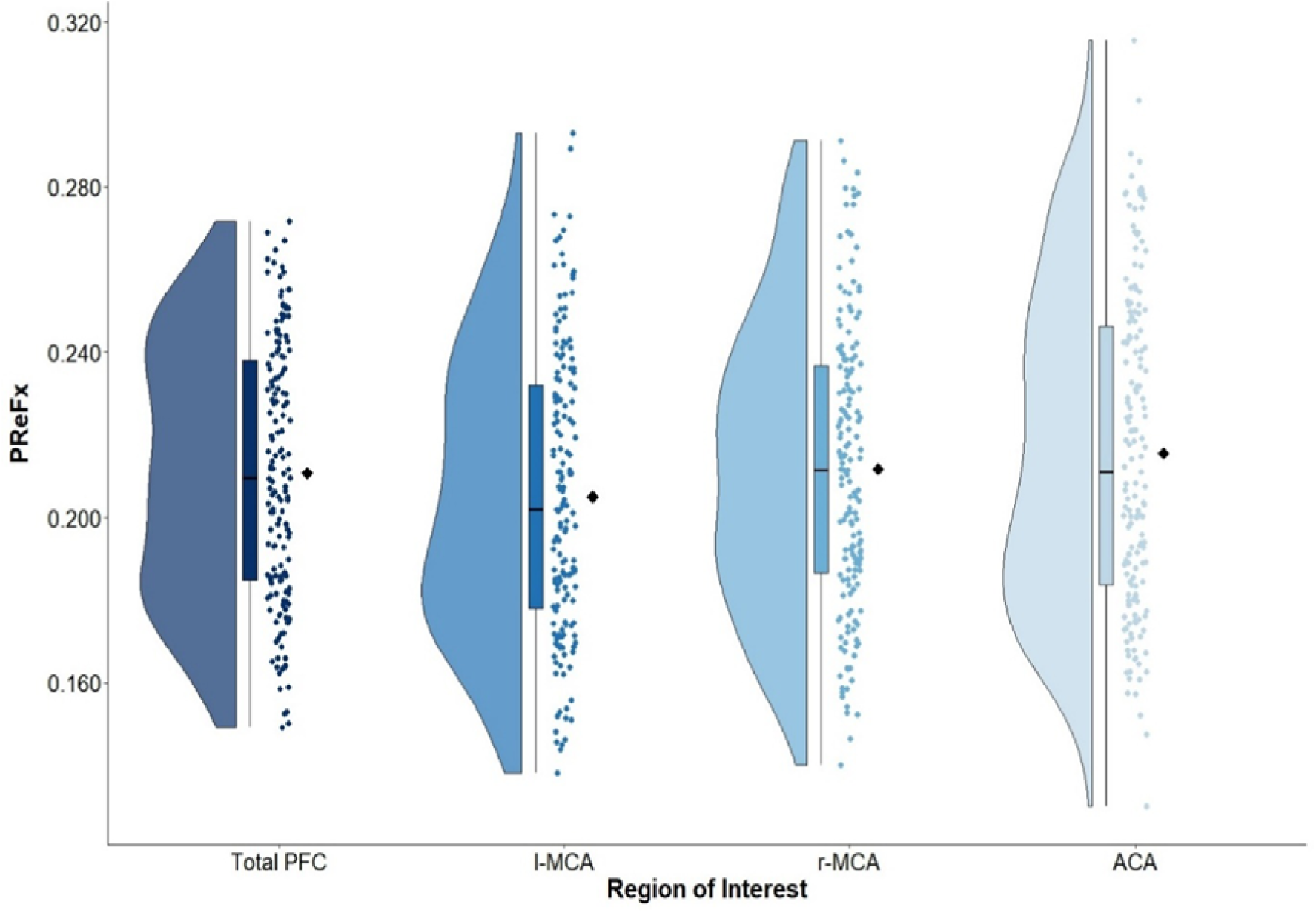
Pulse Relaxation Function (PReFx) at each ROI (Total-PFC, l-MCA, ACA, and r-MCA). Dots represent PReFx value for each individual averaged over all values at that ROI. Black diamonds represent the group mean. Also shown is distribution of scores and boxplots with median (line), interquartile range (box) and range (whiskers). Visualised using ‘ggrain’ (Judd et al., 2024).

#### 3.1.4. Cerebral arterial elasticity – Relationships with age, cognition and cardiovascular health

Table 3B shows associations between PReFx at each ROI and age, cognition, and cardiovascular measures. As the pattern was largely consistent between the three lateralised ROIs and the Total-PFC, here we briefly summarise findings for the Total-PFC.

Higher PReFx was significantly associated with lower age, even within this narrow age range, but not with global cognition score (ACE-III). Higher PReFx was also associated with lower sBP, pulse pressure, heart rate, cholesterol ratio, triglyceride and glucose levels, and higher HDL cholesterol. It was also linked to higher cardiorespiratory fitness and MVPA and lower BMI. Thus, greater cerebral arterial elasticity was associated with younger age, better cardiovascular health profile, and healthier lifestyle indicators.

#### 3.1.5. CVRF Burden on Cerebral Arterial Elasticity

Figure 3 shows average PReFx at the Total-PFC as a function of CVRF Burden group. There was a significant main effect of CVRF Burden on PReFx (F(3, 151) = 5.54, p ≤ .001), with a significant linear trend (F(1,151)=15.65, p ≤ 0.001), indicating that higher CVRF burden was associated with lower arterial elasticity. The effect of CVRF burden did not vary across ROIs (F(6, 302) = 1.51, p = .175).

**Figure 3.**
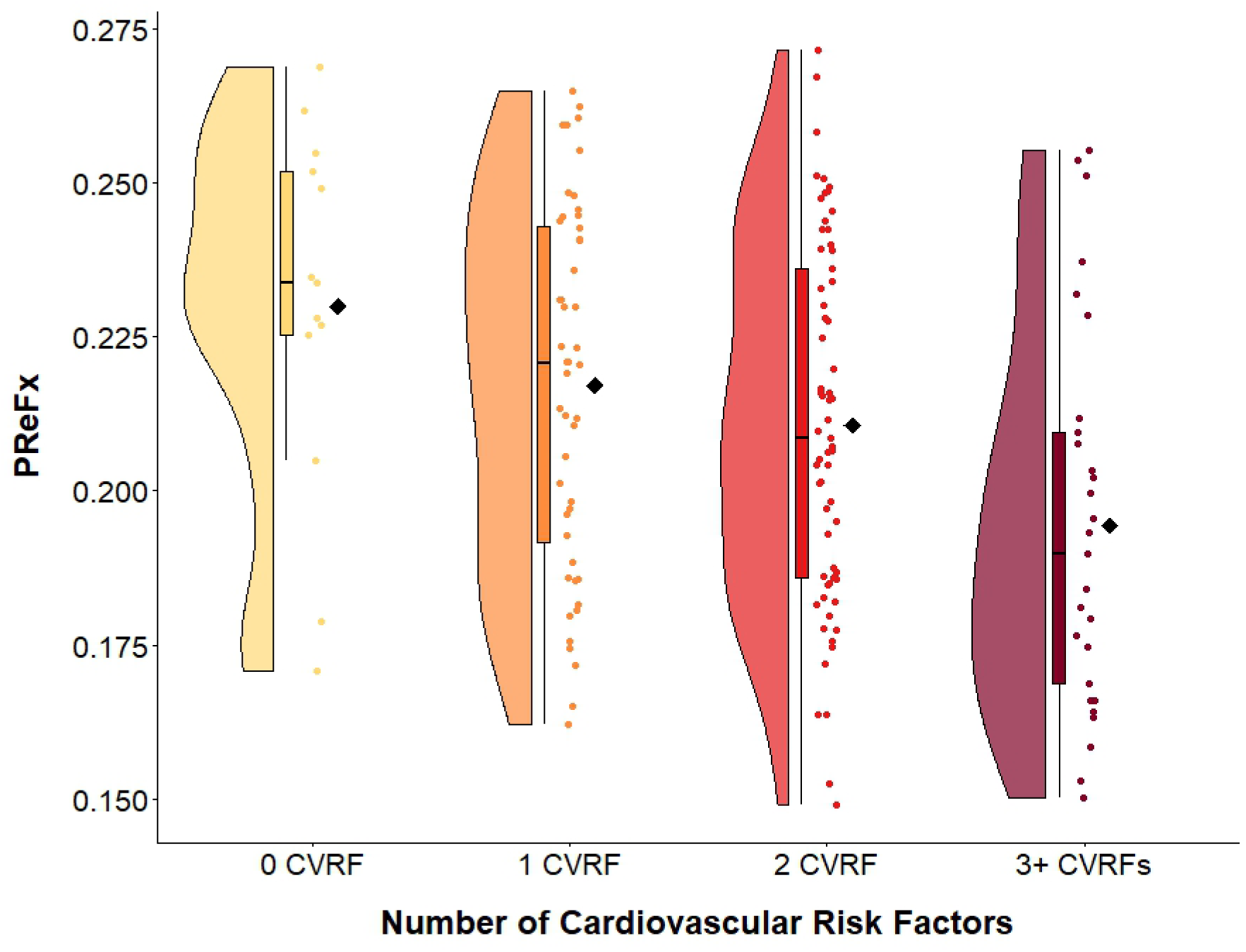
PReFx values at Total-PFC for each CVRF Burden group. Dots represent PReFx value for each individual averaged over all values at that ROI. Black diamonds represent the group mean. Also shown is distribution of scores and boxplots with median (line), interquartile range (box) and range (whiskers). Visualised using ‘ggrain’ (Judd et al., 2024).

Figure 4 compares PReFx maps at each ROI (l-MCA, ACA, r-MCA) for No CVRF and 3+ CVRF groups. At all ROIs, the No CVRF group had mid-high range PReFx values (indicated by yellow), whereas the 3+ CVRF group had notably smaller PReFx values (indicated by red), representing lower arterial elasticity.

**Figure 4.**
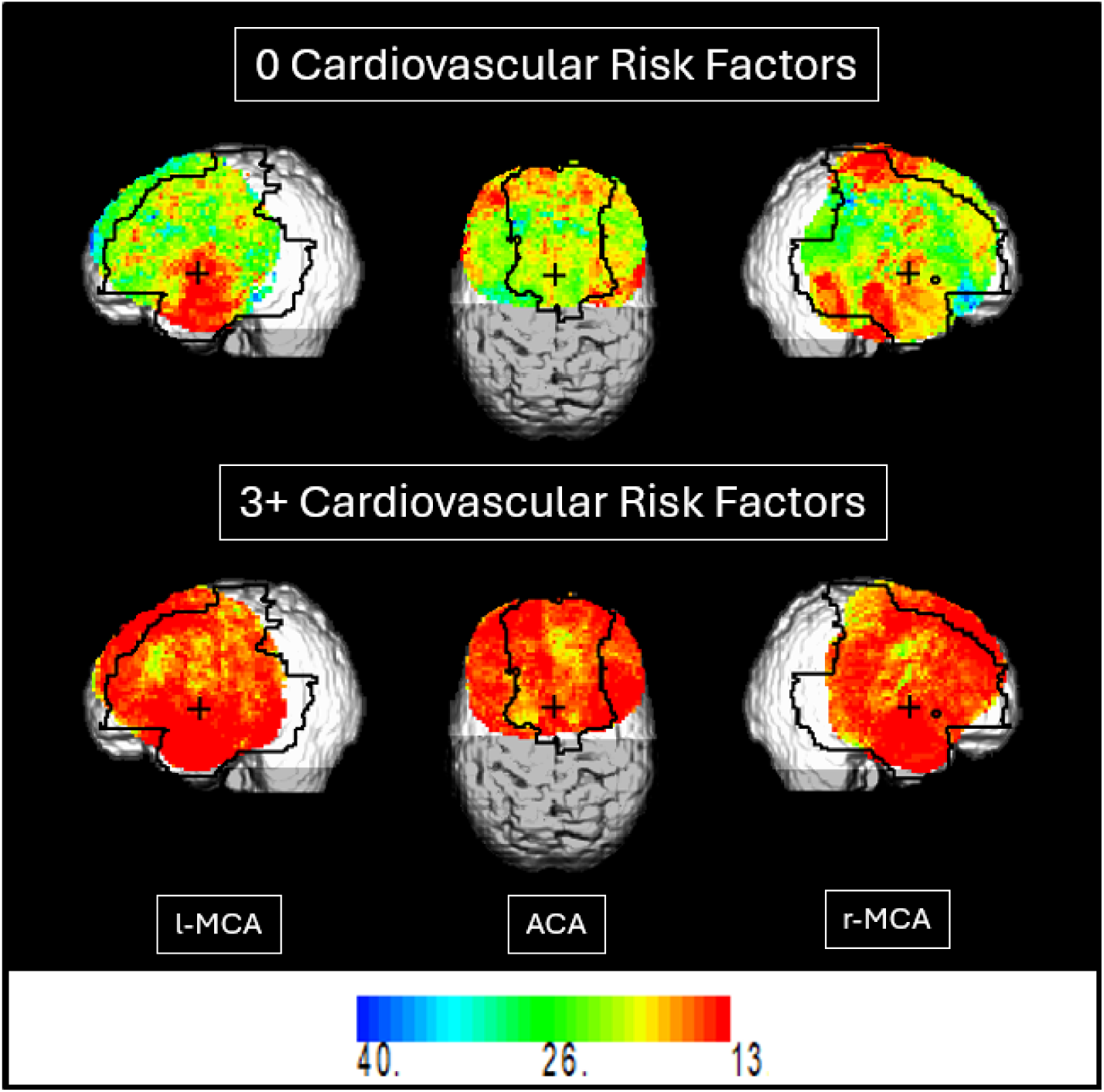
Cerebral arterial elasticity (PReFx) from each ROI covering anterior brain areas perfused by l-MCA, ACA and r-MCA in the 0 CVRF Group (top) and 3+ CVRF Group (bottom). Colder colours represent high PReFx values or more elastic arteries; hotter colours represent lower PReFx values or stiffer arteries).

#### 3.1.6. CVRF mediation of the relationship between age and cerebral arterial elasticity

Given that the effect of age or CVRF burden on PReFx did not differ by ROI, in the following analyses, mediation effects were examined for the average ROI (Total-PFC; Figure 5).

**Figure 5.**
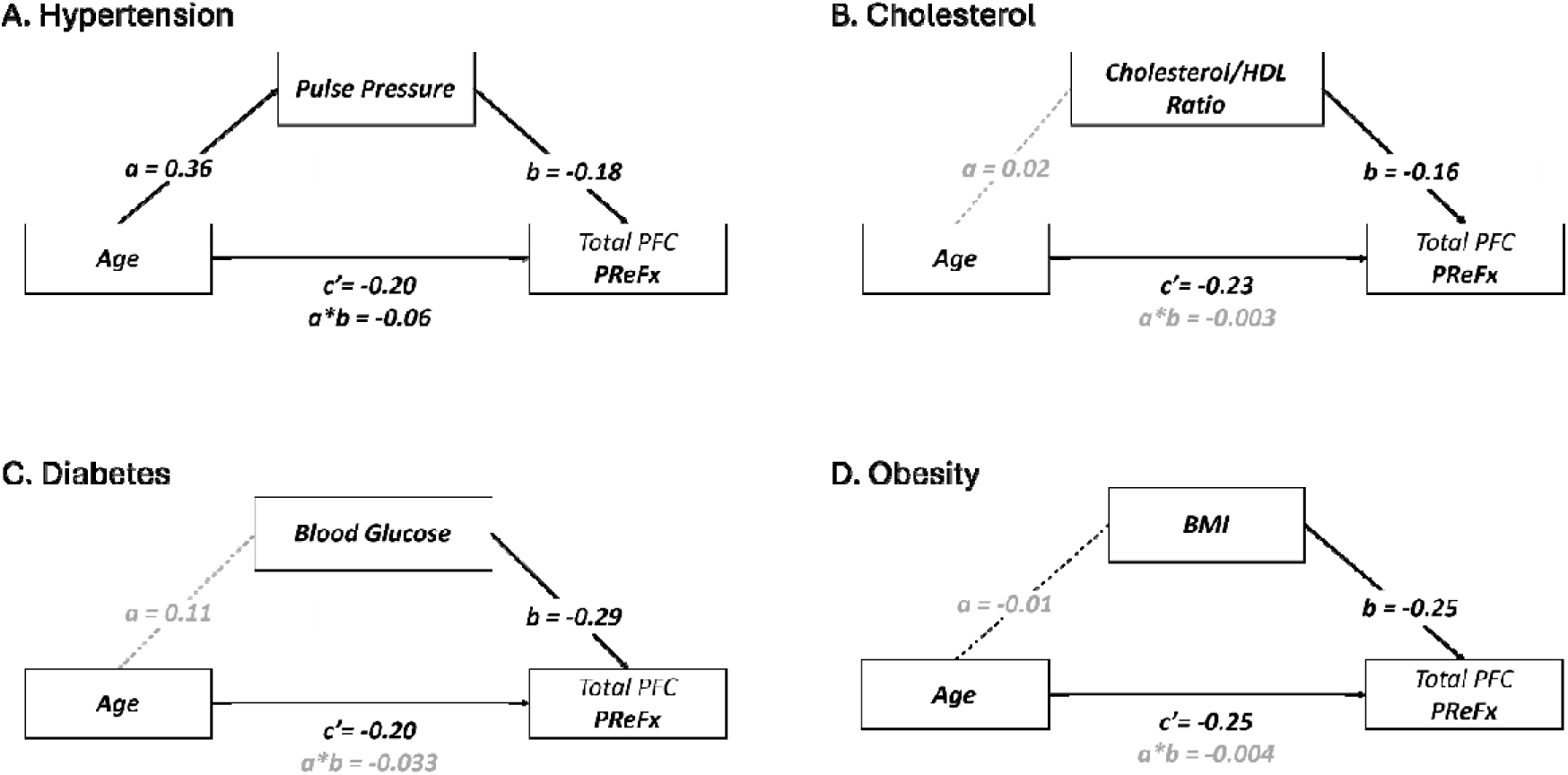
Mediation models testing cardiovascular markers as mediators of the Age-PReFx relationship.

##### Pulse pressure (Figure 5A)

The mediation model was statistically significant (R² = .096, F(2, 156) = 8.26, p < .001), with age and pulse pressure explaining 9.6% of variability in PReFx.

Increasing age was associated with higher pulse pressure (a path: β = 0.36 [.22, .51], p < .001), which in turn was associated with lower PReFx (b path: β = −0.18 [-.34, −.02], p = .030). The direct effect of age on PReFx remained significant when including pulse pressure as a mediator (c’ path: β = −0.20 [-.36, −.03], p = .018). The indirect effect was small but significant (a*b path: β = −0.06 [-.13, −.001]). Thus, pulse pressure partially mediated the relationship between age and PReFx, with the indirect pathway accounting for approximately 24.9% of the total age effect.

##### Cholesterol Ratio (Figure 5B)

The mediation model examining cholesterol ratio was statistically significant (R² = .079, F(2, 155) = 6.61, p = .002). Age and cholesterol ratio explained 7.9% of variability in PReFx. Age was not significantly associated with cholesterol ratio (a path: β = .02 [-.14, .17], p = .847). However, higher cholesterol ratio was associated with lower PReFx (b path: β = −.16 [-.31, −.01], p = .042). The direct effect of age on PReFx remained significant (c’ path: β = −.23 [-.39, −.08], p = .003) when controlling for cholesterol ratio, but the indirect effect was not significant (β = −.003 [-.04, .03]), indicating that cholesterol ratio did not mediate the relationship between age and PReFx.

##### Glucose (Figure 5C)

The mediation model examining glucose was statistically significant (R² = .139, F(2, 155) = 12.48, p < .001). Age and glucose explained 13.9% of variability in PReFx. Age was not significantly associated with glucose (a path: β = .11 [-.05, .27], p = .168). However, higher glucose was associated with lower PReFx (b path: β = −.29 [-.44, −.15], p < .001). The direct effect of age on PReFx remained significant (c’ path: β = −.20 [-.35, −.05], p = .009) when controlling for glucose, but the indirect effect was not significant (β = −.033 [-.09, .02]), indicating that glucose did not mediate the relationship between age and PReFx.

##### BMI (Figure 5D)

The mediation model examining BMI was statistically significant (R² = .125, F(2, 158) = 11.28, p < .001). Age and BMI explained 12.5% of variability in PReFx. Age was not significantly associated with BMI (a path: β = −.01 [-.17, .14], p = .864). However, higher BMI was associated with lower PReFx (b path: β = −.25 [-.40, −.11], p = .001). The direct effect of age on PReFx remained significant (c’ path: β = −.25 [-.39, −.10], p = .001) when controlling for BMI, but the indirect effect was not significant (β = .004 [-.03, .05]), indicating that BMI did not mediate the relationship between age and PReFx.

### 3.2. Longitudinal analyses: Phase 1 (Baseline) vs Phase 2 (1.5 years)

#### 3.2.1. Demographic and cardiovascular indices – change over time

Table 4 shows age, cognitive and cardiovascular data for the sixty-four participants with pulse-DOT data at both phases^5^. Cognition score did not reduce over time, consistent with evidence that ACE-III is stable in healthy controls over 2.5 years (Carrick et al., 2025).

**Table 4.** Change in Cognition and Cardiovascular Health from Phase 1 to Phase 2 (n=64 58% female). Note that fasted bloods and MRI were only available at Phase 1.

| Variable | Phase 1 | Phase 2 | *t* | *df* | *p* |
| --- | --- | --- | --- | --- | --- |
|  | *M (SD)* | *M (SD)* |  |  |  |
| *Age (yrs)* | 65.21 (3.12) | 66.83 (3.11) |  |  |  |
| *ACE-III Total* | 94.07 (4.01) | 94.89 (2.97) | -1.83 | 60 | .071 |
| *Body Mass Index (BMI)* | 27.10 (4.35) | 27.52 (5.85) | -1.12 | 62 | .269 |
| *Systolic Blood Pressure (sBP; mmHg)* | 142.44 (16.48) | 133.88 (13.70) | 4.58 | 62 | <.001 |
| *Diastolic Blood Pressure (dBP; mmHg)* | 78.64 (8.56) | 75.69 (7.70) | 2.91 | 62 | .005 |
| *Pulse Pressure (PP; mmHg)* | 63.81 (12.87) | 58.19 (10.59) | 4.15 | 62 | <.001 |
| *Heart rate (beats/min)* | 64.44 (11.02) | 68.18 (10.97) | -3.82 | 62 | <.001 |

BMI did not change significantly, while heart rate showed a small but significant increase. In contrast, unexpectedly, blood pressure measures significantly *improved* over this interval. Of the 49 people who had been diagnosed with hypertension at Baseline, only 3 (<1%) started taking medication before Follow-up, so change in medication status is unlikely to fully explain this decline. However, medication dosage adjustments may also have occurred for other participants.

#### 3.2.2. Cerebral Arterial Elasticity – change over time

Consistent with Phase 1 data, Phase 2 PReFx scores showed high reliability across resting-state blocks and across regions.

On average, PReFx declined from Phase 1 to Phase 2 by approximately 9% (Figure 6A; *F*(1,63) = 20.12, *p* < .001, η²p = .242). The level of decline varied across ROIs (*F*(2,126) = 11.78, *p* <.001, η²p = .158). Both ACA and r-MCA ROIs showed substantial decline in PReFx over the 1.5 year period (ACA: 10% decline, t(63)=4.01, p<.001, 95%CI [.011 .031]; r-MCA: 11% decline, t(63)=4.11, p<.001, 95%CI [.012 .034]). There was a small, non-significant decline at l-MCA (3.1%; t(63)=1.61, p=.113, 95%CI [-.002 .034]). As l-MCA had lower values at baseline, this could partially reflect a floor effect.

**Figure 6.**
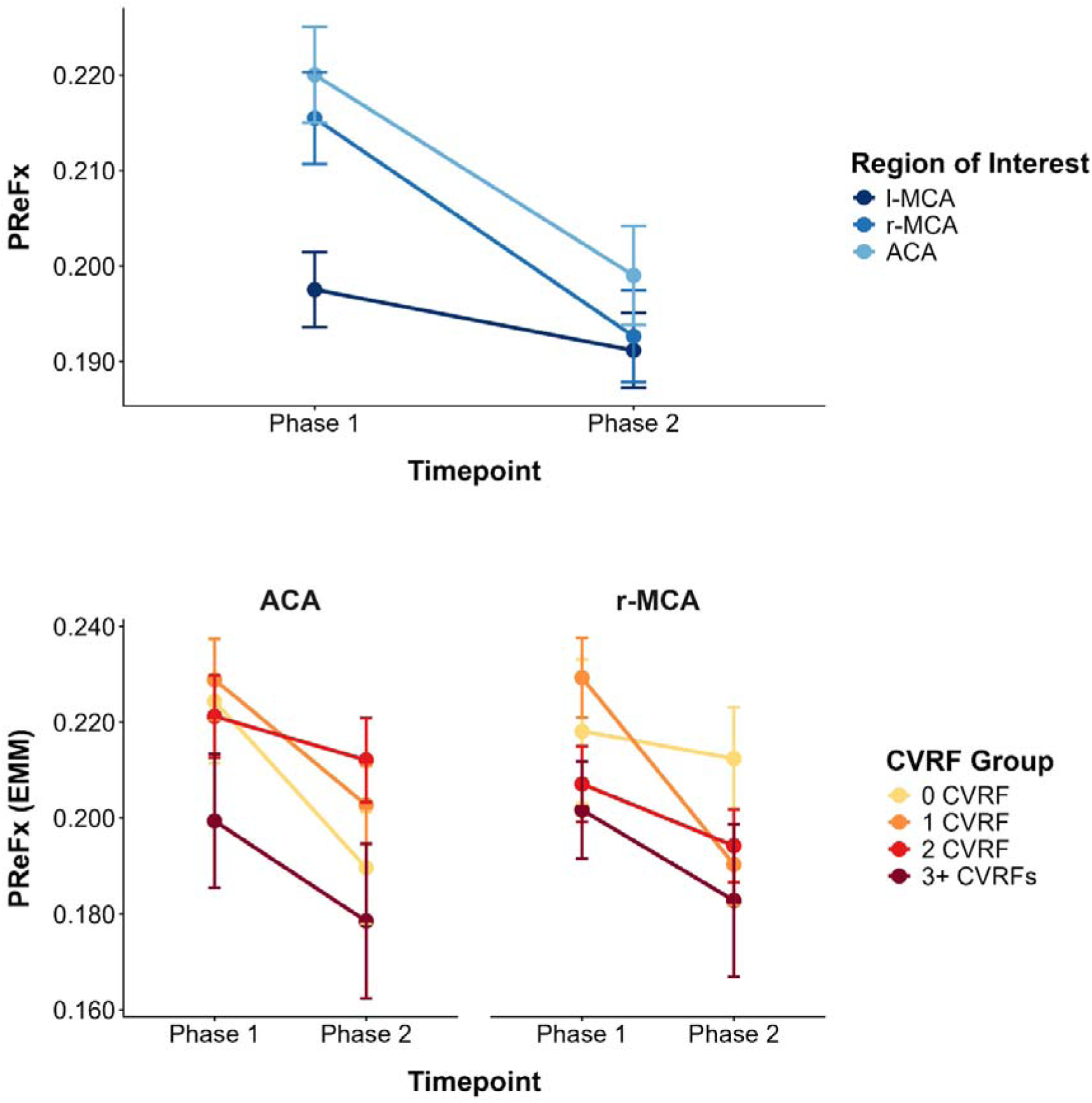
Top Panel: PReFx by Phase and at each ROI. Bottom Panel: PReFx from ACA and r-MCA at each Phase by CVRF Burden. Estimated marginal means with standard error.

Phase 1 and 2 PReFx values were negatively correlated, such that individuals with higher PReFx at Baseline values declined more at Follow-up (*β* = −.549, *p* < .001), a pattern consistent with regression to the mean. However, given the restricted PReFx scale, it is also likely that individuals with higher initial PReFx values had more room for decline.

The level of PReFx decline did not vary between CVRF Burden groups at either ACA or r-MCA (Figure 6B; F(3,57)=1.29, p=.285; F(3,57)=1.52, p=.220). While PReFx decline was positively correlated with age and baseline level of key cardiovascular risk factors, none of these correlations were statistically significant (all p > 0.05).

## 4. Discussion

In this study, we investigated the relationships between cerebral arterial elasticity and cardiovascular health in an age-restricted sample of older adults. Cerebral arterial elasticity was measured over the anterior brain, including the frontal lobes and anterior regions of temporal and parietal lobes, using the pulse relaxation function (PReFx), a metric derived from the arterial pulse wave that is recorded with diffuse optical tomography (Pulse-DOT; Fabiani et al., 2014).

Consistent with previous studies by Fabiani and colleagues (Fabiani et al., 2014; Tan et al., 2019, 2017), PReFx had strong test-retest reliability, with highly consistent values across the two resting state blocks and three ROIs perfused by the anterior and middle cerebral arteries. Nevertheless, PReFx did vary across the anterior brain, with lower arterial elasticity in the left ROI that is perfused by the left MCA compared to both the ACA or right MCA. This is consistent with evidence that left MCA is more sensitive to stroke (Hedna et al., 2013)^6^. Despite a restricted age range (60-70yrs at baseline), PReFx declined with increasing age and was not associated with global cognitive ability, in this cognitively healthy sample. Together, these findings support PReFx as a reliable measure with potential as an early biomarker for cerebral vascular health, particularly in arteries that perfuse the frontal lobes.

### 4.1. Relationship between cerebral arterial elasticity and indices of cardiovascular health

We present the first evidence that cerebral arterial elasticity over anterior brain areas perfused by MCA and ACA varies as a function of cardiovascular risk factor burden in cognitively healthy and physically active older adults. Specifically, increasing number of CVRFs present was linearly associated with lower cerebral arterial elasticity.

Mohammadi et al. (2021) reported similar findings using pulse-DOT measures to examine the association between cerebral arterial health and cardiovascular health status. Specifically, pulse amplitude across frontal and motor cortices was higher in participants with coronary artery disease (CAD), compared to two age-matched control groups, one with low and the other with high Framingham Risk Scores (FRS). Across groups, higher FRS was significantly associated with higher pulse amplitude. Higher pulse amplitude is indicative of less elastic arterial walls and associated with peripheral pulse pressure. In contrast, the groups did not differ in PRF, a measure akin to PReFx used here. Differences in sample size, cohort characteristics (i.e., the Mohammadi study recruited participants with at least one CVRF), and measurement approach may account for these discrepant findings. However, importantly these studies provide converging evidence that poorer cardiovascular health is associated with lower arterial elasticity across frontal brain areas. This is consistent with chronic exposure to higher arterial pulsatility and stiffness in cerebral vessels contributing to microvascular damage (Bateman et al., 2008).

The relationship between increasing CVRF burden and lower cerebral arterial elasticity is also consistent with evidence that cumulative CVRF burden is associated with increased arterial stiffening, in both the aorta (cf-PWV; AlGhatrif et al., 2013; McEniery et al., 2010; Mitchell et al., 2011) and large cerebral arteries (mean flow velocity; Pase et al., 2012). Scuteri et al. (2014) showed that multiple CVRF contribute to aortic stiffness beyond what would be expected from each factor alone. Moreover, specific clusters of risk factors, including impaired glucose tolerance, and elevated blood pressure and triglycerides (with or without abdominal obesity) were associated with greater aortic arterial thickening and stiffening in older adults (Scuteri et al., 2014).

Of the four CVRF metrics, only pulse pressure significantly partially mediated the relationship between age and cerebral arterial elasticity. This is consistent with well-established evidence for blood pressure as a primary determinant of aortic arterial stiffness (AlGhatrif and Lakatta., 2015; Jennings et al., 2020; Safar et al., 2006). Age-related changes to central arteries also impact cerebral arterial health (see Reeve et al., 2024 for review). For example, adults with hypertension have higher pulsatility index in the lenticulostriate arteries (i.e., smaller arteries branching from MCA measured with phase contrast MRI) than those without (van den Kerkhof et al., 2023). Similarly, higher systolic blood pressure is related to increased blood flow velocity in the MCAs (Arts et al., 2022; Zhang et al., 2006).

It is important to note that despite being, on average, very physically active (i.e., average MVPA exceeded Australian recommendations for this age group), this cohort had average systolic blood pressure well above the ‘normal’ level (120 mm Hg) and entering the stage 2 hypertension cutoff (140 mm Hg), with diastolic blood pressure just below the elevated level cutoff (80 mm Hg). This resulted in an average pulse pressure of 77 mm Hg, above the healthy range of 40-60 mg Hg. Moreover, we used current Australian guidelines to define cut-offs for defining presence or absence of hypertension (i.e., 140/90 mmHg; Nelson et al., 2024), which are well above the current US guidelines (130/80 mmHg).

### 4.2. Longitudinal decline in cerebral arterial elasticity

This study provides the first evidence of longitudinal decline in PReFx in cognitively healthy older adults over a 1.5 year interval, highlighting the sensitivity of PReFx to subtle changes in cerebral arterial elasticity. This finding is consistent with evidence that the health of the large cerebral arteries declines over similar timeframes using flow-based measures. For example, in healthy older adults (aged 50+ years), Miller et al. (2021) reported that pulsatility in the MCA and vertebral arteries increased over 17 months, and baseline pulsatility was related to lower fluid cognitive abilities at follow-up. Han et al. (2022) also showed that cerebral blood flow (CBF) declined over 2.1 years in cognitively healthy older adults.

This decline in cerebral arterial elasticity did not significantly vary with level of CVRF burden at baseline. Similarly, the level of individual CVRF at baseline (i.e., cholesterol ratio, pulse pressure, glucose, BMI) was not associated with level of change in arterial elasticity over 1.5 years. It is possible that the relatively short follow-up period may not have been sufficient to capture the effect of these CVRFs on remodelling of cerebral arteries, a process that typically unfolds over many years (Nilsson, 2015). In studies measuring systemic arterial elasticity, elevated systolic blood pressure has been shown to predict increased cf-PWV over 9 years (AlGhatrif et al., 2013) and 16 years (Tomiyama et al., 2023). While some studies have reported accelerated arterial stiffening with CVRF burden over shorter periods (i.e., < 6 years; Benetos et al., 2002; Crichton et al., 2014; Safar et al., 2006), these timeframes still well exceeded the 1.5 year retest interval used here.

### 4.3. Conclusion

In conclusion, we provide the first evidence that cerebral arterial elasticity reduced linearly with increasing CVRF burden and reduced in as little as 1.5 years, in the highly active, cognitively healthy and age-restricted older adult ACTIVate cohort. We also show, for the first time, that brachial pulse pressure partially mediated the relationship between age and cerebral arterial elasticity, consistent with hypertension as a key contributor to arterial stiffness. In contrast, measures of other CVRF (i.e., total cholesterol/HDL ratio, BMI and glucose did not, possibly because they are associated with different cardiovascular and cerebrovascular mechanisms. Moreover, in this relatively small longitudinal sample, neither CVRF burden nor the level of key measures of these CVRF were associated with the rate of decline in arterial elasticity.

Further work is needed in larger samples and over longer periods to replicate these findings and examine mechanisms of action of different CVRFs, perhaps also using other measures of CVRF indicators. It is also necessary to examine interactive effects of medication status and type. This was beyond the scope of the present study, given recruitment was based on convenience sampling and medication status was recorded but could not be systematically controlled.

Nevertheless, the fact that there was evidence of significant decline over 1.5yrs highlights the potential utility of PReFx as a sensitive biomarker for monitoring cerebrovascular ageing and aligns with recent calls for improved vascular biomarkers in dementia research (Sweeney et al., 2019), particularly those that can detect changes before cognitive symptoms emerge.

## Acknowledgements

We thank Felicity Simpson, Nathan Tran, Alexandra Wade, Maddison Mellow, Louise Massie, Kate Dyer, Karen Wilson, Gemma Mieko Furuhashi, Helen Nicholas, Riley Jackson, Teigan Cotterill, Mahmoud Abdolhoseini and Fayeem Aziz for valued contribution towards data collection and study coordination. We also thank the HMRI Research Volunteer Register for partial recruitment of participants at the Newcastle site.

The ACTIVate Study is funded by an NHMRC Boosting Dementia Research Priority Round 5 grant (GNT1171313 to AES (Principal), FK, MF, GG). AES was supported by a Henry Brodaty Dementia Australia Research Foundation mid-career fellowship. MF and GG were funded by NIA grants R01AG059878 and RF1AG062666. JJ, NW were supported by University of Newcastle Postgraduate Research Scholarships awarded by UON for ARC-DP200101471 (FK).

## Data Availability Statement

Due to the ongoing longitudinal nature of the ACTIVate study, the dataset analysed during this study is not publicly available. However, it may be obtained upon reasonable request via the ACTIVate Data Request Form: https://research.unisa.edu.au/redcap/surveys/?s=MEHLHR3TKKX9XMML

## Footnotes

1 The 1.5yr follow-up session only included ACTIVate primary outcome measures. Optical imaging was optional.

2 All participants with dBP>90 also had sBP>140.

3 All participants identified as obese using BMI also had WHR values near or above the established threshold for central adiposity.

4 This participant was retained in subsequent analyses as they had met MoCA screening criteria. Analyses were re-run after removing this participant and produced highly consistent results.

5 Bloods and MRI were not collected at Phase 2.

6 Although this may partly due to left-sided strokes producing symptoms that are more readily detectable (e.g., language and speech impairments (Portegies et al., 2015)).

## References

AlGhatrif, M., Lakatta, E.G., 2015. The conundrum of arterial stiffness, elevated blood pressure, and aging. Current Hypertension Reports 17, 12–12. 10.1007/s11906-014-0523-z

AlGhatrif, M., Strait, J.B., Morrell, C.H., Canepa, M., Wright, J., Elango, P., Scuteri, A., Najjar, S.S., Ferrucci, L., Lakatta, E.G., 2013. Longitudinal trajectories of arterial stiffness and the role of blood pressure: the Baltimore Longitudinal Study of Aging. Hypertension 62, 934–41. 10.1161/HYPERTENSIONAHA.113.01445

Alvarez-Bueno, C., Cunha, P.G., Martinez-Vizcaino, V., Pozuelo-Carrascosa, D.P., Visier-Alfonso, M.E., Jimenez-Lopez, E., Cavero-Redondo, I., 2020. Arterial Stiffness and Cognition Among Adults: A Systematic Review and Meta-Analysis of Observational and Longitudinal Studies. J Am Heart Assoc 9, e014621. 10.1161/JAHA.119.014621

Arts, T., Onkenhout, L.P., Amier, R.P., van der Geest, R., van Harten, T., Kappelle, J., Kuipers, S., van Osch, M.J.P., van Bavel, E.T., Biessels, G.J., Zwanenburg, J.J.M., Consortium, H.-B.C., 2022. Non-Invasive Assessment of Damping of Blood Flow Velocity Pulsatility in Cerebral Arteries With MRI. Journal of Magnetic Resonance Imaging 55, 1785–1794. 10.1002/jmri.27989

Badji, A., Sabra, D., Bherer, L., Cohen-Adad, J., Girouard, H., Gauthier, C.J., 2019. Arterial stiffness and brain integrity: A review of MRI findings. Ageing Res Rev 53, 100907. 10.1016/j.arr.2019.05.001

Bateman, G.A., Levi, C.R., Schofield, P., Wang, Y., Lovett, E.C., 2008. The venous manifestations of pulse wave encephalopathy: windkessel dysfunction in normal aging and senile dementia. Neuroradiology 50, 491–497. 10.1007/s00234-008-0374-x

Bazalar-Palacios, J., Jaime Miranda, J., Carrillo-Larco, R.M., Gilman, R.H., Smeeth, L., Bernabe-Ortiz, A., 2021. Aggregation and combination of cardiovascular risk factors and their association with 10-year all-cause mortality: the PERU MIGRANT Study. BMC Cardiovasc Disord 21, 582. 10.1186/s12872-021-02405-8

Benetos, A., Adamopoulos, C., Bureau, J.M., Temmar, M., Labat, C., Bean, K., Thomas, F., Pannier, B., Asmar, R., Zureik, M., Safar, M., Guize, L., 2002. Determinants of accelerated progression of arterial stiffness in normotensive subjects and in treated hypertensive subjects over a 6-year period. Circulation 105, 1202–7. 10.1161/hc1002.105135

Benetos, A., Safar, M., Rudnichi, A., Smulyan, H., Richard, J.-L., Ducimetière, P., Guize, L., 1997. Pulse Pressure. Hypertension 30, 1410–1415. 10.1161/01.HYP.30.6.1410

Benjamini, Y., Hochberg, Y., 1995. Controlling the False Discovery Rate: A Practical and Powerful Approach to Multiple Testing. Journal of the Royal Statistical Society. Series B (Methodological) 57, 289–300.

Bowie, D.C., Low, K.A., Rubenstein, S.L., Islam, S.S., Zimmerman, B., Camacho, P.B., Sutton, B.P., Gratton, G., Fabiani, M., 2024. Neurovascular mechanisms of cognitive aging: Sex-related differences in the average progression of arteriosclerosis, white matter atrophy, and cognitive decline. Neurobiology of Disease 201, 106653. 10.1016/j.nbd.2024.106653

Brunner, E.J., Shipley, M.J., Ahmadi-Abhari, S., Tabak, A.G., McEniery, C.M., Wilkinson, I.B., Marmot, M.G., Singh-Manoux, A., Kivimaki, M., 2015. Adiposity, Obesity, and Arterial Aging: Longitudinal Study of Aortic Stiffness in the Whitehall II Cohort. Hypertension 66, 294–300. 10.1161/HYPERTENSIONAHA.115.05494

Bustamante Gallo, J. P., del Neglia Cermeño, C.S., Díaz-Ortega, J.L., Yupari-Azabache, I.L., 2024. Predictive Models of Atherogenic Risk in Citizens of Trujillo (Peru) Based on Associated Factors. Nutrients 16, 4138. 10.3390/nu16234138

Cao, X., Zhang, L., Wang, X., Chen, Z., Zheng, C., Chen, L., Zhou, H., Cai, J., Hu, Z., Tian, Y., Gu, R., Huang, Y., Wang, Z., 2023. Cardiovascular disease and all-cause mortality associated with individual and combined cardiometabolic risk factors. BMC Public Health 23, 1725. 10.1186/s12889-023-16659-8

Carrick, J., Cheung, S.C., Foxe, D., Piguet, O., 2025. Interpreting Addenbrooke’s Cognitive Examination-III Scores in Dementia: Performance Distributions and Clinically Meaningful Change. Euro J of Neurology 32, e70257. 10.1111/ene.70257

Castelli, W.P., Garrison, R.J., Wilson, P.W., Abbott, R.D., Kalousdian, S., Kannel, W.B., 1986. Incidence of coronary heart disease and lipoprotein cholesterol levels. The Framingham Study. JAMA 256, 2835–2838.

Cecelja, M., Chowienczyk, P., 2013. Arterial stiffening: Causes and consequences. Artery Research 7, 22–27. 10.1016/j.artres.2012.09.001

Chen, H., Chen, Y., Wu, W., Cai, Z., Chen, Z., Yan, X., Wu, S., 2021. Total cholesterol, arterial stiffness, and systolic blood pressure: a mediation analysis. Sci Rep 11, 1330. 10.1038/s41598-020-79368-x

Chiarelli, A.M., Fletcher, M.A., Tan, C.H., Low, K.A., Maclin, E.L., Zimmerman, B., Kong, T., Gorsuch, A., Gratton, G., Fabiani, M., 2017. Individual differences in regional cortical volumes across the life span are associated with regional optical measures of arterial elasticity. NeuroImage 162, 199–213. 10.1016/j.neuroimage.2017.08.064

Chiarelli, A.M., Maclin, E.L., Fabiani, M., Gratton, G., 2015. A kurtosis-based wavelet algorithm for motion artifact correction of fNIRS data. NeuroImage 112, 128–137. 10.1016/j.neuroimage.2015.02.057

Cohen, J.B., Mitchell, G.F., Gill, D., Burgess, S., Rahman, M., Hanff, T.C., Ramachandran, V.S., Mutalik, K.M., Townsend, R.R., Chirinos, J.A., 2022. Arterial Stiffness and Diabetes Risk in Framingham Heart Study and UK Biobank. Circulation Research 131, 545–554. 10.1161/CIRCRESAHA.122.320796

Crichton, Georgina E., Elias, M.F., Davey, A., Sullivan, K.J., Robbins, M.A., 2014. Higher HDL cholesterol is associated with better cognitive function: the Maine-Syracuse study. J Int Neuropsychol Soc 20, 961–970. 10.1017/S1355617714000885

Crichton, G. E., Elias, M.F., Robbins, M.A., 2014. Cardiovascular health and arterial stiffness: the Maine-Syracuse Longitudinal Study. J Hum Hypertens 28, 444–449. 10.1038/jhh.2013.131

Elias, M.F., Crichton, G.E., Dearborn, P.J., Robbins, M.A., Abhayaratna, W.P., 2017. Associations between Type 2 Diabetes Mellitus and Arterial Stiffness: A Prospective Analysis Based on the Maine-Syracuse Study. Pulse 5, 88–98. 10.1159/000479560

Elias, M.F., Robbins, M.A., Budge, M.M., Abhayaratna, W.P., Dore, G.A., Elias, P.K., 2009. Arterial Pulse Wave Velocity and Cognition With Advancing Age. Hypertension 53, 668–673. 10.1161/HYPERTENSIONAHA.108.126342

Fabiani, M., Low, K.A., Tan, C., Zimmerman, B., Fletcher, M.A., Schneider-Garces, N., Maclin, E.L., Chiarelli, A.M., Sutton, B.P., Gratton, G., 2014a. Taking the pulse of aging: Mapping pulse pressure and elasticity in cerebral arteries with optical methods. Psychophysiology 51, 1072–1088. 10.1111/psyp.12288

Farnsworth Von Cederwald, B., Josefsson, M., Wåhlin, A., Nyberg, L., Karalija, N., 2022. Association of Cardiovascular Risk Trajectory With Cognitive Decline and Incident Dementia. Neurology 98, e2013–e2022. 10.1212/WNL.0000000000200255

Feng, S., Zeng, F.-A., Chance, B., 1995. Photon migration in the presence of a single defect: a perturbation analysis. Applied Optics 34, 3826. 10.1364/AO.34.003826,

Fujita, S., Mori, S., Onda, K., Hanaoka, S., Nomura, Y., Nakao, T., Yoshikawa, T., Takao, H., Hayashi, N., Abe, O., 2023. Characterization of Brain Volume Changes in Aging Individuals With Normal Cognition Using Serial Magnetic Resonance Imaging. JAMA Netw Open 6, e2318153. 10.1001/jamanetworkopen.2023.18153

Gratton, G., 2000. “Opt-cont” and “Opt-3D”: A software suite for the analysis and 3D reconstruction of the event-related optical signal (EROS), in: Psychophysiology. CAMBRIDGE UNIV PRESS 40 WEST 20TH STREET, NEW YORK, NY 10011-4211 USA, pp. S44–S44.

Grover, S.A., Palmer, C.S., Coupal, L., 1994. Serum Lipid Screening to Identify High-Risk Individuals for Coronary Death: The Results of the Lipid Research Clinics Prevalence Cohort. Arch Intern Med 154, 679–684. 10.1001/archinte.1994.00420060113012

Haider, A.W., Larson, M.G., Franklin, S.S., Levy, D., 2003. Systolic Blood Pressure, Diastolic Blood Pressure, and Pulse Pressure as Predictors of Risk for Congestive Heart Failure in the Framingham Heart Study. Ann Intern Med 138, 10–16. 10.7326/0003-4819-138-1-200301070-00006

Hajjar, I., Goldstein, F.C., Martin, G.S., Quyyumi, A.A., 2016. Roles of Arterial Stiffness and Blood Pressure in Hypertension-Associated Cognitive Decline in Healthy Adults. Hypertension 67, 171–175. 10.1161/HYPERTENSIONAHA.115.06277

Han, H., Lin, Z., Soldan, A., Pettigrew, C., Betz, J.F., Oishi, K., Li, Y., Liu, P., Albert, M., Lu, H., 2022. Longitudinal Changes in Global Cerebral Blood Flow in Cognitively Normal Older Adults: A Phase-Contrast MRI Study. J Magn Reson Imaging 56, 1538–1545. 10.1002/jmri.28133

Hayes, A.F., 2022. Introduction to mediation, moderation, and conditional process analysis: a regression-based approach, Third edition. ed, Methodology in the social sciences. Guilford Publications, New York.

Hedna, V.S., Bodhit, A.N., Ansari, S., Falchook, A.D., Stead, L., Heilman, K.M., Waters, M.F., 2013. Hemispheric Differences in Ischemic Stroke: Is Left-Hemisphere Stroke More Common? J Clin Neurol 9, 97–102. 10.3988/jcn.2013.9.2.97

Herzog, M.J., Müller, P., Lechner, K., Stiebler, M., Arndt, P., Kunz, M., Ahrens, D., Schmeißer, A., Schreiber, S., Braun-Dullaeus, R.C., 2025. Arterial stiffness and vascular aging: mechanisms, prevention, and therapy. Sig Transduct Target Ther 10, 282. 10.1038/s41392-025-02346-0

Hsieh, S., Schubert, S., Hoon, C., Mioshi, E., Hodges, J.R., 2013. Validation of the Addenbrooke’s Cognitive Examination III in frontotemporal dementia and Alzheimer’s disease. Dement Geriatr Cogn Disord 36, 242–250. 10.1159/000351671

Ikonomidis, I., Thymis, J., 2023. The vicious circle of arterial elasticity, blood pressure, glycemia, and renal function. Hypertens Res 46, 1599–1602. 10.1038/s41440-023-01262-6

Jennings, J.R., Muldoon, M.F., Allen, B., Ginty, A.T., Gianaros, P.J., 2020. Cerebrovascular function in hypertension: Does high blood pressure make you old? Psychophysiology. 10.1111/psyp.13654

Jurca, R., Jackson, A.S., LaMonte, M.J., Morrow, J.R., Blair, S.N., Wareham, N.J., Haskell, W.L., Van Mechelen, W., Church, T.S., Jakicic, J.M., Laukkanen, R., 2005. Assessing Cardiorespiratory Fitness Without Performing Exercise Testing. American Journal of Preventive Medicine 29, 185–193. 10.1016/j.amepre.2005.06.004

Kaess, B.M., Rong, J., Larson, M.G., Hamburg, N.M., Vita, J.A., Levy, D., Benjamin, E.J., Vasan, R.S., Mitchell, G.F., 2012. Aortic stiffness, blood pressure progression, and incident hypertension. JAMA 308, 875–881. 10.1001/2012.jama.10503

Kim, Young-Joo, Kim, Yun-Jin, Cho, B.-M., Lee, S., 2010. Metabolic syndrome and arterial pulse wave velocity. Acta Cardiol 65, 315–321. 10.2143/AC.65.3.2050348

Kohn, J.C., Lampi, M.C., Reinhart-King, C.A., 2015. Age-related vascular stiffening: causes and consequences. Front. Genet. 06. 10.3389/fgene.2015.00112

Kong, T.S., Gratton, C., Low, K.A., Tan, C.H., Chiarelli, A.M., Fletcher, M.A., Zimmerman, B., Maclin, E.L., Sutton, B.P., Gratton, G., Fabiani, M., 2020. Age-related differences in functional brain network segregation are consistent with a cascade of cerebrovascular, structural, and cognitive effects. Network Neuroscience 4, 89–114. 10.1162/netn_a_00110

Kovács, B., Cseprekál, O., Diószegi, Á., Lengyel, S., Maroda, L., Paragh, G., Harangi, M., Páll, D., 2022. The Importance of Arterial Stiffness Assessment in Patients with Familial Hypercholesterolemia. J Clin Med 11, 2872. 10.3390/jcm11102872

LaPlume, A.A., McKetton, L., Levine, B., Troyer, A.K., Anderson, N.D., 2022. The adverse effect of modifiable dementia risk factors on cognition amplifies across the adult lifespan. Alzheimers Dement (Amst) 14, e12337. 10.1002/dad2.12337

Liu, Q., Fang, J., Cui, C., Dong, S., Gao, L., Bao, J., Li, Y., Ma, M., Chen, N., He, L., 2021. Association of Aortic Stiffness and Cognitive Decline: A Systematic Review and Meta-Analysis. Front Aging Neurosci 13, 680205. 10.3389/fnagi.2021.680205

Mailey, E.L., White, S.M., Wójcicki, T.R., Szabo, A.N., Kramer, A.F., McAuley, E., 2010. Construct validation of a non-exercise measure of cardiorespiratory fitness in older adults. BMC Public Health 10, 59. 10.1186/1471-2458-10-59

Mancusi, C., Losi, M.A., Izzo, R., Canciello, G., Carlino, M.V., Albano, G., De Luca, N., Trimarco, B., de Simone, G., 2018. Higher pulse pressure and risk for cardiovascular events in patients with essential hypertension: The Campania Salute Network. Eur J Prev Cardiol 25, 235–243. 10.1177/2047487317747498

McEniery, C.M., Spratt, M., Munnery, M., Yarnell, J., Lowe, G.D., Rumley, A., Gallacher, J., Ben-Shlomo, Y., Cockcroft, J.R., Wilkinson, I.B., 2010. An analysis of prospective risk factors for aortic stiffness in men: 20-year follow-up from the Caerphilly prospective study. Hypertension 56, 36–43. 10.1161/HYPERTENSIONAHA.110.150896

Mellow, M.L., Dumuid, D., Olds, T., Stanford, T., Dorrian, J., Wade, A.T., Fripp, J., Xia, Y., Goldsworthy, M.R., Karayanidis, F., Breakspear, M.J., Smith, A.E., 2024. Cross-sectional associations between 24-hour time-use composition, grey matter volume and cognitive function in healthy older adults. Int J Behav Nutr Phys Act 21, 11. 10.1186/s12966-023-01557-4

Mellow, M.L., Dumuid, D., Wade, A.T., Stanford, T., Olds, T.S., Karayanidis, F., Hunter, M., Keage, H.A.D., Dorrian, J., Goldsworthy, M.R., Smith, A.E., 2022. Twenty-four-hour time-use composition and cognitive function in older adults: cross-sectional findings of the ACTIVate study. Front Hum Neurosci 16, 1051793. 10.3389/fnhum.2022.1051793

Millán, J., Pintó, X., Muñoz, A., Zúñiga, M., Rubiés-Prat, J., Pallardo, L.F., Masana, L., Mangas, A., Hernández-Mijares, A., González-Santos, P., Ascaso, J.F., Pedro-Botet, J., 2009. Lipoprotein ratios: Physiological significance and clinical usefulness in cardiovascular prevention. Vasc Health Risk Manag 5, 757–765.

Miller, M.L., Ghisletta, P., Jacobs, B.S., Dahle, C.L., Raz, N., 2021. Changes in cerebral arterial pulsatility and hippocampal volume: a transcranial doppler ultrasonography study. Neurobiology of Aging 108, 110–121. 10.1016/j.neurobiolaging.2021.08.014

Mitchell, G.F., Van Buchem, M.A., Sigurdsson, S., Gotal, J.D., Jonsdottir, M.K., Kjartansson, Ó., Garcia, M., Aspelund, T., Harris, T.B., Gudnason, V., Launer, L.J., 2011. Arterial stiffness, pressure and flow pulsatility and brain structure and function: the Age, Gene/Environment Susceptibility – Reykjavik Study. Brain 134, 3398–3407. 10.1093/brain/awr253

Mohammadi, H., Vincent, T., Peng, K., Nigam, A., Gayda, M., Fraser, S., Joanette, Y., Lesage, F., Bherer, L., 2021. Coronary artery disease and its impact on the pulsatile brain: A functional NIRS study. Human Brain Mapping 42, 3760–3776. 10.1002/hbm.25463

Muhammad, I.F., Borné, Y., Östling, G., Kennbäck, C., Gottsäter, M., Persson, M., Nilsson, P.M., Engström, G., 2017. Arterial Stiffness and Incidence of Diabetes: A Population-Based Cohort Study. Diabetes Care 40, 1739–1745. 10.2337/dc17-1071

Nabeel, P.M., Chandran, D.S., Kaur, P., Thanikachalam, S., Sivaprakasam, M., Joseph, J., 2021. Association of incremental pulse wave velocity with cardiometabolic risk factors. Sci Rep 11, 15413. 10.1038/s41598-021-94723-2

Najjar, S.S., Scuteri, A., Lakatta, E.G., 2005. Arterial aging: Is it an immutable cardiovascular risk factor? Hypertension 46, 454–462. 10.1161/01.HYP.0000177474.06749.98

Najjar, S.S., Scuteri, A., Shetty, V., Wright, J.G., Muller, D.C., Fleg, J.L., Spurgeon, H.P., Ferrucci, L., Lakatta, E.G., 2008. Pulse wave velocity is an independent predictor of the longitudinal increase in systolic blood pressure and of incident hypertension in the Baltimore Longitudinal Study of Aging. J Am Coll Cardiol 51, 1377–1383. 10.1016/j.jacc.2007.10.065

Nelson, M.R., Banks, E., Brown, A., Chow, C.K., Peiris, D.P., Stocks, N.P., Davies Ao, R., Raffoul, N., Kalman, L., Bradburn, E., Jennings, G., 2024. 2023 Australian guideline for assessing and managing cardiovascular disease risk. Med J Aust 220, 482–490. 10.5694/mja2.52280

Nilsson, P.M., 2015. The Concept of Early Vascular Ageing – An Update in 2015. EMJ Diabet 80–86. 10.33590/emjdiabet/10312465

Ohyama, Y., Teixido-Tura, G., Ambale-Venkatesh, B., Noda, C., Chugh, A.R., Liu, C.-Y., Redheuil, A., Stacey, R.B., Dietz, H., Gomes, A.S., Prince, M.R., Evangelista, A., Wu, C.O., Hundley, W.G., Bluemke, D.A., Lima, J.A.C., 2016. Ten-year longitudinal change in aortic stiffness assessed by cardiac MRI in the second half of the human lifespan: the multi-ethnic study of atherosclerosis. Eur Heart J Cardiovasc Imaging 17, 1044–1053. 10.1093/ehjci/jev332

O’Rourke, M.F., Hashimoto, J., 2007. Mechanical Factors in Arterial Aging. Journal of the American College of Cardiology 50, 1–13. 10.1016/j.jacc.2006.12.050

Pase, M.P., Grima, N.A., Stough, C.K., Scholey, A., Pipingas, A., 2012. Cardiovascular Disease Risk and Cerebral Blood Flow Velocity. Stroke 43, 2803–2805. 10.1161/STROKEAHA.112.666727

Pase, M.P., Himali, J.J., Mitchell, G.F., Beiser, A., Maillard, P., Tsao, C., Larson, M.G., DeCarli, C., Vasan, R.S., Seshadri, S., 2016. Association of Aortic Stiffness With Cognition and Brain Aging in Young and Middle-Aged Adults: The Framingham Third Generation Cohort Study. Hypertension 67, 513–519. 10.1161/HYPERTENSIONAHA.115.06610

Perdomo, S.J., 2019. Clinical relevance of brachial pulse pressure as a measure of cerebrovascular disease risk. J Clin Hypertens (Greenwich) 21, 1016–1017. 10.1111/jch.13582

Peters, R., Peters, J., Booth, A., Anstey, K.J., 2020. Trajectory of blood pressure, body mass index, cholesterol and incident dementia: systematic review. The British Journal of Psychiatry 216, 16–28. 10.1192/bjp.2019.156

Portegies, M.L.P., Selwaness, M., Hofman, A., Koudstaal, P.J., Vernooij, M.W., Ikram, M.A., 2015. Left-Sided Strokes Are More Often Recognized Than Right-Sided Strokes. Stroke 46, 252–254. 10.1161/STROKEAHA.114.007385

Prenner, S.B., Chirinos, J.A., 2015. Arterial stiffness in diabetes mellitus. Atherosclerosis 238, 370–379. 10.1016/j.atherosclerosis.2014.12.023

R Core Team, 2025. A Language and Environment for Statistical Computing R Foundation for Statistical Computing, Vienna, Austria.

Reeve, E.H., Barnes, J.N., Moir, M.E., Walker, A.E., 2024. Impact of arterial stiffness on cerebrovascular function: a review of evidence from humans and preclincal models. Am J Physiol Heart Circ Physiol 326, H689–H704. 10.1152/ajpheart.00592.2023

Rolls, E.T., Huang, C.-C., Lin, C.-P., Feng, J., Joliot, M., 2020. Automated anatomical labelling atlas 3. Neuroimage 206, 116189. 10.1016/j.neuroimage.2019.116189

Safar, M.E., Thomas, F., Blacher, J., Nzietchueng, R., Bureau, J.-M., Pannier, B., Benetos, A., 2006. Metabolic Syndrome and Age-Related Progression of Aortic Stiffness. Journal of the American College of Cardiology 47, 72–75. 10.1016/j.jacc.2005.08.052

Said, M.A., Eppinga, R.N., Lipsic, E., Verweij, N., Van Der Harst, P., 2018. Relationship of Arterial Stiffness Index and Pulse Pressure With Cardiovascular Disease and Mortality. JAHA 7, e007621. 10.1161/JAHA.117.007621

Scuteri, A., Cunha, P.G., Rosei, E.A., Badariere, J., Bekaert, S., Cockcroft, J.R., Cotter, J., Cucca, F., Buyzere, M.L.D., Meyer, T.D., Ferrucci, L., Franco, O., Gale, N., Gillebert, T.C., Hofman, A., Langlois, M., Laucevicius, A., Laurent, S., Raso, F.U.S.M., Morrell, C.H., Muiesan, M.L., Munnery, M.M., Navickas, R., Oliveira, P., Orru’, M., Pilia, M.G., Rietzschel, E.R., Ryliskyte, L., Salvetti, M., Schlessinger, D., Sousa, N., Stefanadis, C., Strait, J., Daele, C.V., Villa, I., Vlachopoulos, C., Witteman, J., Xaplanteris, P., Nilsson, P., Lakatta, E.G., 2014. Arterial stiffness and influences of the metabolic syndrome: A cross-countries study. Atherosclerosis 233, 654–660. 10.1016/j.atherosclerosis.2014.01.041

Scuteri, A., Najjar, S.S., Muller, D.C., Andres, R., Hougaku, H., Metter, E.J., Lakatta, E.G., 2004. Metabolic syndrome amplifies the age-associated increases in vascular thickness and stiffness. J Am Coll Cardiol 43, 1388–1395. 10.1016/j.jacc.2003.10.061

Singer, J., Trollor, J.N., Baune, B.T., Sachdev, P.S., Smith, E., 2014. Arterial stiffness, the brain and cognition: A systematic review. Ageing Research Reviews 15, 16–27. 10.1016/j.arr.2014.02.002

Smith, A.E., Wade, A.T., Olds, T., Dumuid, D., Breakspear, M.J., Laver, K., Goldsworthy, M.R., Ridding, M.C., Fabiani, M., Dorrian, J., Hunter, M., Paton, B., Abdolhoseini, M., Aziz, F., Mellow, M.L., Collins, C., Murphy, K.J., Gratton, G., Keage, H., Smith, R.T., Karayanidis, F., 2022. Characterising activity and diet compositions for dementia prevention: protocol for the ACTIVate prospective longitudinal cohort study. BMJ Open 12, e047888. 10.1136/bmjopen-2020-047888

Sweeney, M.D., Montagne, A., Sagare, A.P., Nation, D.A., Schneider, L.S., Chui, H.C., Harrington, M.G., Pa, J., Law, M., Wang, D.J.J., Jacobs, R.E., Doubal, F.N., Ramirez, J., Black, S.E., Nedergaard, M., Benveniste, H., Dichgans, M., Iadecola, C., Love, S., Bath, P.M., Markus, H.S., Salman, R.A., Allan, S.M., Quinn, T.J., Kalaria, R.N., Werring, D.J., Carare, R.O., Touyz, R.M., Williams, S.C.R., Moskowitz, M.A., Katusic, Z.S., Lutz, S.E., Lazarov, O., Minshall, R.D., Rehman, J., Davis, T.P., Wellington, C.L., González, H.M., Yuan, C., Lockhart, S.N., Hughes, T.M., Chen, C.L.H., Sachdev, P., O’Brien, J.T., Skoog, I., Pantoni, L., Gustafson, D.R., Biessels, G.J., Wallin, A., Smith, E.E., Mok, V., Wong, A., Passmore, P., Barkof, F., Muller, M., Breteler, M.M.B., Román, G.C., Hamel, E., Seshadri, S., Gottesman, R.F., van Buchem, M.A., Arvanitakis, Z., Schneider, J.A., Drewes, L.R., Hachinski, V., Finch, C.E., Toga, A.W., Wardlaw, J.M., Zlokovic, B. V., 2019. Vascular dysfunction—The disregarded partner of Alzheimer’s disease. Alzheimer’s & Dementia 15, 158–167. 10.1016/J.JALZ.2018.07.222

Takase, H., Dohi, Y., Toriyama, T., Okado, T., Tanaka, S., Sonoda, H., Sato, K., Kimura, G., 2011. Brachial-ankle pulse wave velocity predicts increase in blood pressure and onset of hypertension. Am J Hypertens 24, 667–673. 10.1038/ajh.2011.19

Tan, C.H., Low, K.A., Chiarelli, A.M., Fletcher, M.A., Navarra, R., Burzynska, A.Z., Kong, T.S., Zimmerman, B., Maclin, E.L., Sutton, B.P., Gratton, G., Fabiani, M., 2019. Optical measures of cerebral arterial stiffness are associated with white matter signal abnormalities and cognitive performance in normal aging. Neurobiology of Aging 84, 200–207. 10.1016/j.neurobiolaging.2019.08.004

Tan, C.H., Low, K.A., Kong, T., Fletcher, M.A., Zimmerman, B., Maclin, E.L., Chiarelli, A.M., Gratton, G., Fabiani, M., 2017. Mapping cerebral pulse pressure and arterial compliance over the adult lifespan with optical imaging. PLoS ONE 12, e0171305. 10.1371/journal.pone.0171305

Tomiyama, H., Imai, T., Shiina, K., Higashi, Y., Nakano, H., Takahashi, T., Fujii, M., Matsumoto, C., Yamashina, A., Chikamori, T., 2023. Lifelong Heterogeneous Contribution of Cardiovascular Risk Factors to Slow and Fast Progression of Arterial Stiffness. Hypertension 80, 2159–2168. 10.1161/HYPERTENSIONAHA.123.21481

van den Kerkhof, M., van der Thiel, M.M., van Oostenbrugge, R.J., Postma, A.A., Kroon, A.A., Backes, W.H., Jansen, J.F., 2023. Impaired damping of cerebral blood flow velocity pulsatility is associated with the number of perivascular spaces as measured with 7T MRI. J Cereb Blood Flow Metab 43, 937–946. 10.1177/0271678X231153374

Wilson, J., Webb, A.J.S., 2020. Systolic Blood Pressure and Longitudinal Progression of Arterial Stiffness: A Quantitative Meta-Analysis. Journal of the American Heart Association 9, e017804. 10.1161/JAHA.120.017804

Wittich, W., Phillips, N., Nasreddine, Z.S., Chertkow, H., 2010. Sensitivity and Specificity of the Montreal Cognitive Assessment Modified for Individuals who are Visually Impaired. Journal of Visual Impairment & Blindness 104, 360–368. 10.1177/0145482X1010400606

Yekutieli, D., Benjamini, Y., 1999. Resampling-based false discovery rate controlling multiple test procedures for correlated test statistics. Journal of Statistical Planning and Inference 82, 171–196. 10.1016/S0378-3758(99)00041-5

Zaninotto, P., Batty, G.D., Allerhand, M., Deary, I.J., 2018. Cognitive function trajectories and their determinants in older people: 8 years of follow-up in the English Longitudinal Study of Ageing. J Epidemiol Community Health 72, 685–694. 10.1136/jech-2017-210116

Zhang, P., Huang, Y., Li, Y., Lu, M., Wu, Y., 2006. A large-scale study on relationship between cerebral blood flow velocity and blood pressure in a natural population. J Hum Hypertens 20, 742–748. 10.1038/sj.jhh.1002068

Zheng, M., Zhang, X., Chen, S., Song, Y., Zhao, Q., Gao, X., Wu, S., 2020. Arterial Stiffness Preceding Diabetes. Circulation Research 127, 1491–1498. 10.1161/CIRCRESAHA.120.317950

Zhou, Y., Xu, H.-L., Lin, X.-L., Chen, Z.-T., Ye, Q.-Y., Zhao, Z.-H., 2025. The nonlinear association of ratio of total cholesterol to high density lipoprotein with cognition ability: evidence from a community cohort in China. Front. Nutr. 12. 10.3389/fnut.2025.1525348

Zimmerman, B., Rypma, B., Gratton, G., Fabiani, M., 2021. Age-related changes in cerebrovascular health and their effects on neural function and cognition: A comprehensive review. Psychophysiology 58, e13796. 10.1111/psyp.13796

